# Combining Sampling Methods with Attractor Dynamics in Spiking Models of Head-Direction Systems

**DOI:** 10.1101/2025.02.25.640158

**Authors:** Vojko Pjanovic, Jacob Zavatone-Veth, Paul Masset, Sander Keemink, Michele Nardin

## Abstract

Uncertainty is a fundamental aspect of the natural environment, requiring the brain to infer and integrate noisy signals to guide behavior effectively. Sampling-based inference has been proposed as a mechanism for dealing with uncertainty, particularly in early sensory processing. However, it is unclear how to reconcile sampling-based methods with operational principles of higher-order brain areas, such as attractor dynamics of persistent neural representations. In this study, we present a spiking neural network model for the head-direction (HD) system that combines sampling-based inference with attractor dynamics. To achieve this, we derive the required spiking neural network dynamics and interactions to perform sampling from a large family of probability distributions—including variables encoded with Poisson noise. We then propose a method that allows the network to update its estimate of the current head direction by integrating angular velocity samples—derived from noisy inputs—with a pull towards a circular manifold, thereby maintaining consistent attractor dynamics. This model makes specific, testable predictions about the HD system that can be examined in future neurophysiological experiments: it predicts correlated subthreshold voltage fluctuations; distinctive short- and long-term firing correlations among neurons; and characteristic statistics of the movement of the neural activity “bump” representing the head direction. Overall, our approach extends previous theories on probabilistic sampling with spiking neurons, offers a novel perspective on the computations responsible for orientation and navigation, and supports the hypothesis that sampling-based methods can be combined with attractor dynamics to provide a viable framework for studying neural dynamics across the brain.

## Introduction

Biological organisms live in a world fraught with uncertainty. To survive, they must interpret and act upon uncertain information [5, 6, 7, 8]. This challenge spans multiple levels of neural processing, from low-level sensory processing to complex cognitive decision-making [9, 10]. In particular, noise in the encoding of behaviorally relevant variables can increase the risk of erroneous downstream decisions if not properly managed [11]. The head direction (HD) system is a prominent example of a high-level computation dealing with uncertainty: to keep track of the heading direction, noisily encoded rotational velocity estimates need to be inferred and integrated into a circular attractor [12, 4, 13, 14] (Fig. 1A).

**Figure 1.**
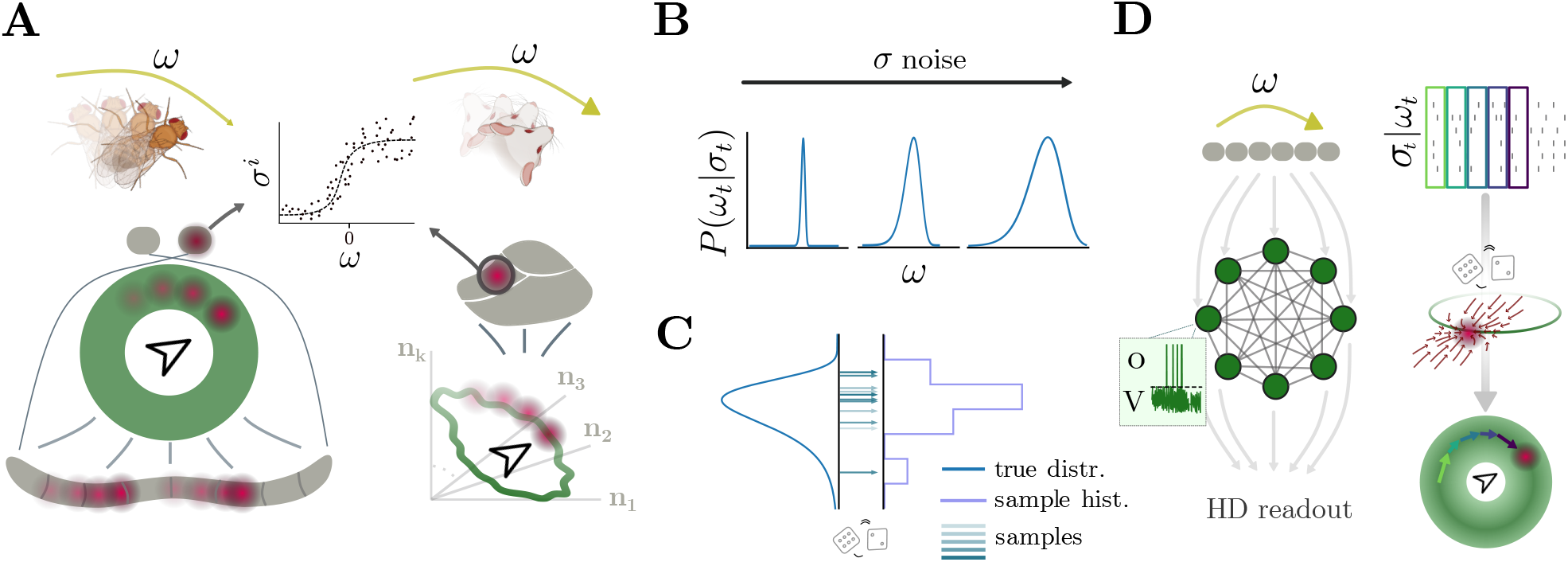
Head direction networks receive noisy measurements from which the head direction must be inferred. **(A)** Angular velocity *ω* is encoded noisily by segregated populations of neurons in different species. Left: GLNO cells in fruit flies are the sole source of rotational velocity information to PEN neurons in the PB [1], which is bidirectionally connected to the EPG neurons in the EB — cartoon depicts simplified anatomy: two round structures depict noduli (NO), grey handlebar-shaped structure depicts PB, and green circle depicts EB; red blob depicts activity. Right: brainstem’s dorsal tegmental nucleus (DTN) cells in rats provide angular velocity information to lateral mammillary nuclei (LMN) neurons [2], whose activity lies on a circular manifold in a high−*D* space [3] and propagate HD information to the rest of the brain [4] — grey cartoon depicts brainstem, red blob activity. Central inset shows cartoon tuning function for one velocity-encoding cell *σ*^*i*^. **(B)** For increasing noise levels on the encoding of the angular velocity, the posterior distribution *P* (*ω*_*t*_|***σ***_*t*_) will show more and more uncertainty. **(C)** Estimating true density and statistics through sampling (colorful arrows: samples, grey line: samples histogram, blue line: true distribution). **(D)** Spike Coding Networks (SCNs) can formalize computations with spikes. Top: angular velocity inputs to SCN, encoded as noisy Poisson spikes. Middle: SCN cartoon that implements angular velocity sampling and HD integration and attraction; inset shows the activity of a single neuron: *V* is the voltage, which depends on external inputs and recurrent interactions Eq. (1); When *V > T*, the neuron emits spikes *o*. Bottom: HD readout is updated according to the sampled angular velocities. The left column illustrates the network structure, the right column the Poisson inputs, the computations of the network, and the resulting read-out.

The HD system is observed in a wide range of species, including vertebrates like rodents, bats, primates, birds and fish [15, 16, 17, 18, 19], and invertebrates, such as locusts and fruit flies [20, 21]. Empirical evidence indicates that angular velocity signals are encoded noisily [2, 22, 1]. Noise makes it impossible to recover the true latent variable; from a theoretical perspective, the best one can do is to consider a posterior distribution of possible angular velocities (Fig. 1B). Further computations given this distribution, such as integration to keep track of head direction, can pose significant mathematical and neural challenges [23, 24, 7]. Even though the brain, either through evolution or plasticity [25], might have found a way to directly approximate the mean of such posterior and use it to update the current HD representation, we will consider an enticing alternative: integration through posterior sampling (Fig. 1C). In statistical sampling methods, successive samples are drawn from an underlying probability distribution, thereby — using enough samples — approximating the distribution. This has the advantage of preserving properties of the original distribution (such as asymmetry, amount of uncertainty, etc.) while immediately providing a stimulus estimate for downstream areas to use. Sampling-based methods thus offer a potential approach for representing and computing with uncertainty in neural systems, and have accumulated substantial experimental support [26, 27, 10, 28, 29].

While previous work mainly focused on sampling-based inference in the early stages of environmental stimulus encoding — with implementations in rate-based [30, 24, 31, 32, 33, 34] and spiking neural networks [35, 36, 37, 38, 39, 40, 41, 42, 43] — it remains unclear how to bridge the statistical inference perspective with the dynamical systems accounts of neural population dynamics. Among these accounts, ring attractor networks have been studied for decades to model continuous attractors in neural circuits, particularly for the HD signals [44, 45, 46, 47, 48, 49, 50]. In classical models, the ring attractor emerges from recurrent connectivity that stabilizes a “bump” of activity encoding a continuous variable such as orientation or direction. Biologically, there is substantial evidence that attractor-like mechanisms are at play in HD systems of rodents [15, 3] and the central complex of the fruit fly [21, 51, 13]. However, many of the classical models consider noiseless angular velocity inputs [44, 47, 48] or, in the case of noisy inputs, do not cast the problem as an inference problem [52] (but see [53]). This gap motivates us to combine the inference-based view with the dynamical properties of ring attractor networks.

We propose a sampling-based HD model that infers angular velocity from noisy inputs and integrates them in a stable fashion by attracting towards a circular manifold. We implement this model using Spike Coding Networks (SCNs) (Fig. 1D) [54, 41, 55, 56, 42], a class of spiking neural networks that allows us to implement computations through dynamics [57]. SCNs are well-suited for implementing the HD system due to their biological realism [58, 55, 59, 60] and fully derivable nature. While previous implementations primarily focused on sampling from Gaussian distributions [41, 42], here we show how to extend these approaches to handle non-Gaussian distributions. This extension enables sampling-based ring attractor dynamics for HD inference and, in turn, clarifies the precise network, cellular, and synaptic requirements for implementing HD computations. Our HD model provides a novel perspective on neural computations involved in orientation and navigation, and offers testable predictions regarding subthreshold voltage correlations, neural activity correlation patterns, and statistics of bump movement.

## Results

To maintain an internal representation of the head direction of an animal, an HD system must overcome several challenges. First, it must infer the head rotation velocity from noisy inputs [2, 22, 1]. Second, it must use the inferred angular velocity and potentially other multimodal inputs to infer and update the current head direction estimate, usually within circularity constraints [21, 3, 50]. Finally, the whole computational process must be possible with spiking neurons, as is clearly the case in vertebrates [4] and likely also in invertebrates [22, 20, 61]. To meet these challenges, we propose a formulation that combines statistical sampling methods, spiking neural network theory, and attractor dynamics. First, we derive how spiking neural networks can perform sampling from arbitrary distributions to accommodate non-Gaussian noisy information. Second, we show how such networks can perform sampling-based inference of stimuli from Poisson neuron inputs. Third, we combine inference, integration, and attractor dynamics in a single HD network. Finally, we provide a list of experimentally testable predictions stemming from this model, as well as the data, experiments, and techniques needed to test these predictions.

### Structural assumptions

HD systems are found in both vertebrates and invertebrates [15, 16, 17, 18, 19, 20, 21] (Fig. 1A). The precise details of each species can be intricate; for example, in fruit flies, the HD circuit has been characterized thoroughly [62], and the mechanisms that facilitate angular velocity integration have been studied in detail [21, 51, 13, 22, 63, 1]. In our study, we are interested in the high-level general principles guiding angular velocity integration and consider the following generic setup (Fig. 1D). We assume that the current angular velocity is encoded by a population of neurons that emit spikes according to Poisson statistics (Fig. 1D top). A downstream brain area, modeled as a recurrent spiking neural network (SNN), receives these spikes and, through local recurrent dynamics, infers the current angular velocity and integrates it to maintain an HD representation (Fig. 1D middle). The HD is read out by downstream areas (or an experimenter) through a linear readout (Fig. 1D bottom). This setup allows us to derive network dynamics and interactions from theory and to propose novel HD computational principles that might underly real biological networks, with the goal of generating testable predictions to inspire future experiments.

### Langevin sampling in Spiking Neural Networks

To implement HD computations, we must first consider how a SNN can represent arbitrary probability distributions. One approach is to compute certain statistics, such as mean and variance; this approach neglects information about the underlying distribution shape in non-Gaussian cases. Instead, in the sampling approach, random samples are drawn from the distribution to recover the true density or arbitrary statistics, which are guaranteed to converge to their true values given enough samples. This is the process of ‘statistical sampling’ (Fig. 1C, Fig. 2A).

**Figure 2.**
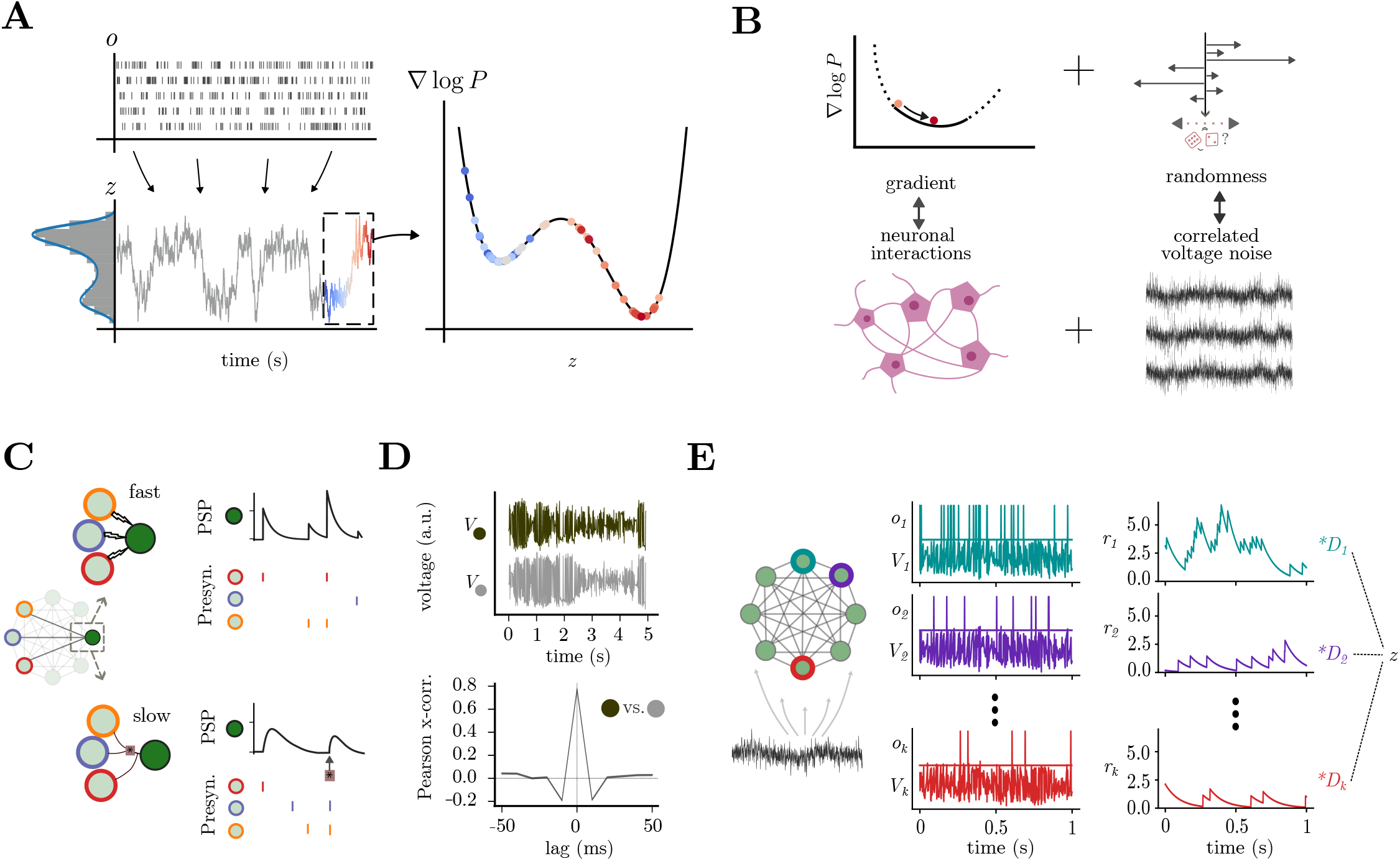
Spiking neural networks can sample from arbitrary distributions through a combination of nonlinear synaptic interactions and correlated voltage fluctuations. **(A)** Example SCN sampling from a given bimodal distribution, a clear example of non-Gaussian distribution, which can be extended to higher orders and higher dimensions(Supp. Fig. 1). Top left: Spikes from the network **o**. Bottom left: network readout *z*_*t*_ corresponding to samples from *P* (*z*). Over time, samples from SCN approximate the original probability distribution, as seen in the histogram/density overlay. Right: the samples **z** explore the gradient landscape of the log *P* distribution; colored dots correspond to samples in the dashed box on the left. **(B)**Langevin dynamics allow one to sample from arbitrary probability distributions. This entails following the slope of the log distribution (∇*logP*) in concert with a source of stochasticity to promote exploration. In our networks, these terms are effectuated by neuronal interactions and correlated voltage stochastic fluctuations, respectively. **(C)** SCNs have two types of interactions between neurons: fast interactions (top) quickly propagate spikes **o** through **Ω**_*f*_, and are responsible for maintaining E/I balance; slow interactions (bottom) act on a slower timescale through **Ω**_*s*_, and implement both linear and nonlinear interactions. Linear interactions alone are sufficient for sampling from Gaussian distributions; generic distributions require nonlinear interactions. **(D)** Correlated voltage stochastic fluctuations enables the network to explore the distribution effectively; the Pearson cross-correlation (bottom) between two voltage traces (top) shows an interesting example of short-timescale subthreshold correlation, reminiscent of experimental studies [64]. **(E)** Left: SCN structure with correlated voltage inputs ***η***. Middle: Voltage *V* and spikes *o* for 3 example neurons; spikes *o* contribute to the post-synaptic potentials *r*, which obey 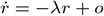. Right: readout **z** can be linearly decoded from the network activity using the decoding vectors *D*_*i*_s: **z** = **Dr**.

A promising approach for performing statistical sampling with spiking neural networks is Langevin sampling, as proposed in previous work [41, 65, 42]. This sampling method uses 1) dynamics, which follow the gradient of a probability distribution, and 2) stochasticity, to explore the entire probability landscape [66] (Fig. 2A,B). Building upon previous studies [41, 56, 42], we show that Langevin sampling from a large family of distributions can be directly implemented by spiking networks with slow, nonlinear interactions of the form

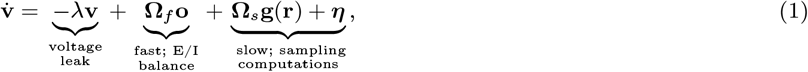

where **v** ∈ ℝ^*N*^ corresponds to neuron voltages, *N* is the number of neurons, *λ* is the voltage leak, **o** ∈ ℝ^*N*^ is the spike vector, and **r** ∈ ℝ^*N*^ are post-synaptic potentials defined as 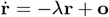. A spike is emitted when the voltage of a neuron crosses a threshold *T*_*i*_; **g**(**r**) ∈ ℝ^*L*^ are sets of possibly nonlinear interactions between neural activities modeling synaptic and/or dendritic interactions (Fig. 2C), and so *L* could be much larger than *N* ; ***η*** ∈ ℝ^*N*^ represents correlated voltage stochastic fluctuations fed into the neuron voltages (Fig. 1D, Fig. 2B-D, Methods). The neurons interact over two timescales:

- Fast timescale interactions: **Ω**_*f*_ ∈ ℝ^*N*×*N*^ is the fast connectivity matrix, acting on spike **o** timescales, through which the network maintains an excitation/inhibition balance [54, 55] (Fig. 2C, top). This timescale is similar to AMPA and GABA_*a*_ receptors, with rapid activation and decay [67].
- Slow timescale interactions: **Ω**_*s*_ ∈ ℝ^*N*×*L*^ is the slow connectivity matrix, mediating slower and potentially nonlinear interactions (**g**(**r**)) (Fig. 2C, bottom). These connections enforce the desired network output dynamics over time [54, 56, 68]. In biology, this timescale is similar to NMDA and GABA_*b*_ receptors, with slow rise and longer-lasting activations [69, 70].

The information sampled by this network is then represented through a linear read-out (Fig. 2E):

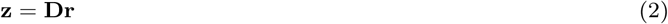

The exact form of the neural interactions **g**(**r**) and the slow weights **Ω**_*s*_ (Methods), discussed in the next section, make sure that **z** evolves according to the gradient of an underlying probability distribution. Finally, the stochastic fluctuations ***η*** drive the network to explore the full distribution (Fig. 2A,B,D).

#### Synaptic interactions required for sampling generic distributions

Previous SCN samplers were limited to Gaussian distributions [41, 42]. Here, we show how to set nonlinear interactions to sample from a wider family of distributions (Eq. (37), Supp. Fig. 1). We achieve this explicitly for exponential family distributions with polynomial moments, for which we derive the exact form and weighting required of the slow, nonlinear interactions **Ω**_*s*_**g**(**r**). Briefly, a polynomial exponential family requires multiplicative dependencies between the variables, which are implemented within the network by multiplicative interactions [56] between the neural activities (Fig. 2B, Supp. Fig. 1), similar to the implementation of nonlinear dynamical systems [56]. Notice that this is not the only option; other synaptic/dendritic nonlinearities can be employed for implementing or approximating different distributions [71, 68].

As an example, the multi-peaked distribution (Fig. 2A) requires interactions of the form *r*_*i*_*r*_*j*_*r*_*k*_, where three pre-synaptic neural activities act multiplicatively at the post-synaptic cell (Fig. 2C). Our formalism allows for sampling distributions in any dimension with arbitrary polynomial moments without the need for approximations (Supp. Fig. 1A). Higher-order polynomials require increasingly complicated multiplicative interactions (Supp. Fig. 1B-C). Specifically, networks have to implement (*g* − 1)-th order multiplications for distributions with *g*-th order polynomial moments (see Methods). For very high orders, such precise interactions become biologically unrealistic, but they can be approximated by combining several lower-order interactions [56].

This approach formalizes precisely the synaptic requirements for implementing Langevin sampling from non-Gaussian distributions in this class of spiking neural networks. Alternative formalisms that can be used to approximate multiplicative interactions, as well as how they are implemented in real biological networks, are summarized in the Discussion.

#### Correlated stochastic fluctuations required for sampling

The voltage stochastic fluctuations ***η*** ensure that the network state drifts and covers the entire space defined by the probability distribution. The dimension and correlation structure of the stochastic fluctuation is determined by the dimensionality of the distribution from which the network is sampling (Methods). From a biological perspective, these sources of stochasticity might arise from external inputs, oscillations, gap junctions, or other sources from the shared extracellular environment [64], and they give rise to correlated subthreshold voltages (Fig. 2C), similar to those found in experiments [72, 73, 74, 75]. Our assumptions give clear functional relevance to these subthreshold correlations, which are otherwise hard to clarify.

#### Quantifying sampling performance

Finally, we quantify how well the network output approximates the desired distribution for different parameter sets (Appendix and Supp. Fig. 2). The sampling performance is affected by various choices of parameters of the networks, such as number of neurons, voltage decay, and readout weights. The network outputs quickly converge to the desired distribution for a large range of parameter sets. Our basic implementation could be extended to improve convergence speed [33, 42]; nonetheless, it provides solid performance for the inference of dynamic stimuli, as we will see in the next section. All in all, our work clarifies how several types of interactions over different timescales, together with voltage stochastic fluctuation, could come together to perform network-level sampling from a very general class of distributions.

### Sampling-based inference of angular velocity from Poisson neuron inputs

The HD system receives noisily encoded angular velocity inputs (Fig. 1, Fig. 3A, left), and in this section, we will focus on building a network that receives the noisy inputs and outputs angular velocity stimulus estimates. Though we focus on circular velocity variables, the same procedure works for generic variables (Methods). Specifically, sampling-based inference consists of drawing samples from the posterior distribution over noisily encoded variables (Fig. 3A, right). This has two main advantages, as it allows downstream circuits to 1) take into account the full uncertainty for further computation and 2) act fast by immediately using the samples received [43, 10]. Moreover, sampling-based inference can help simplify various problems, such as nonlinear state evaluation or prediction of the future [7, 24], and it can be parallelized easily [76, 41].

**Figure 3.**
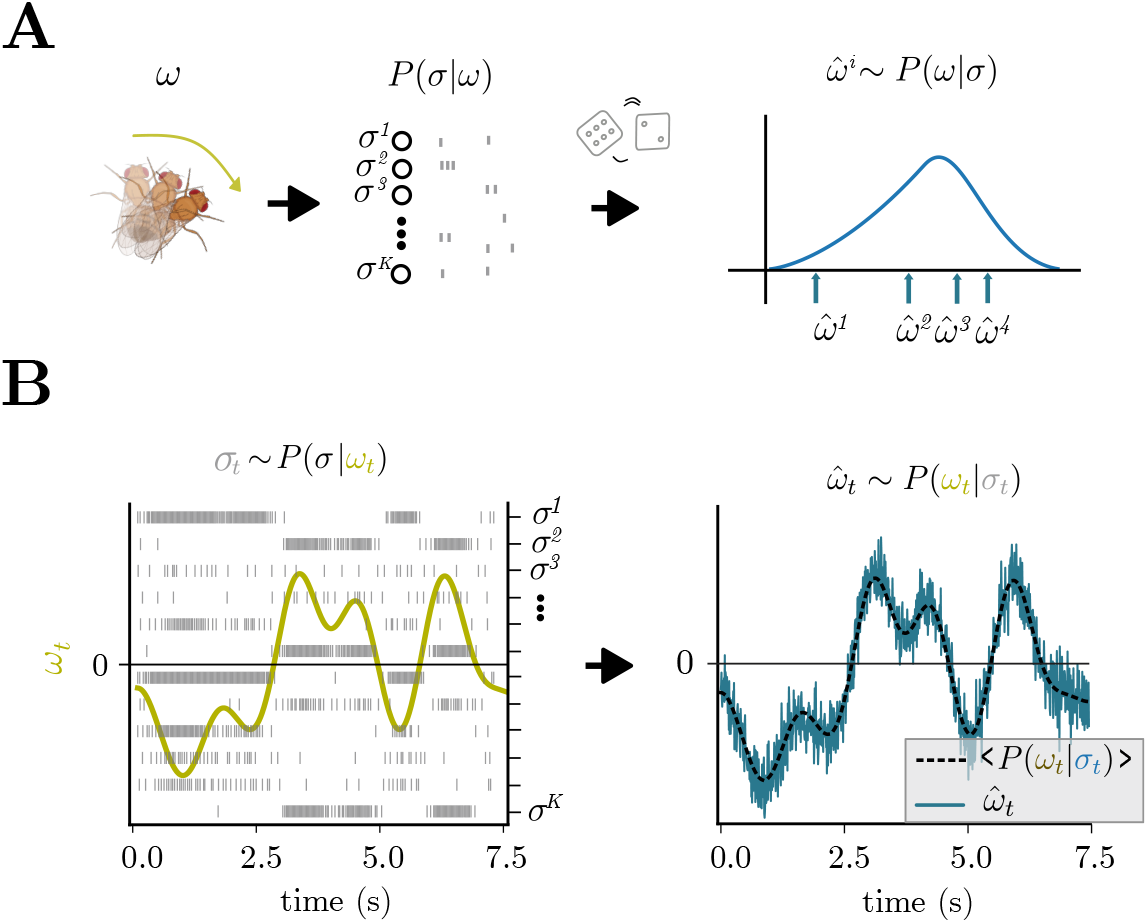
Sampling-based inference of Poisson encoded angular velocity signals. **(A)** Angular velocity is encoded by noisy Poisson neurons; sampling-based inference entails drawing samples from the posterior distribution over angular velocity. **(B)** SCN of the form Eq. (48) can perform sampling based inference of dynamically encoded angular velocity. Left: grey spikes 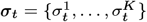 encode a time-varying angular velocity signal *ω*_*t*_. Right: The network readout, 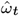, draws samples from the time-varying posterior distribution *P* (*ω*_*t*_|***σ***_*t*_).

We assume that a population of Poisson neurons 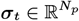 encodes the current angular velocity *ω*_*t*_ ∈ ℝ (Fig. 1A) (See Methods for different firing rate kernels and encoding noise models). The network needs to sample from the posterior distribution over *ω*_*t*_ given the input ***σ***_*t*_:

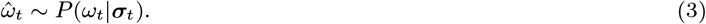

Sampling from this posterior can be formally implemented through Langevin dynamics (Methods) and implemented in an SCN using Eq. (1), as it was done previously for Gaussian variables [41, 42]. Explicitly, this results in a network of the form:

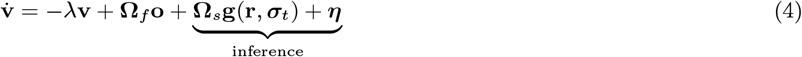

This equation resembles Eq. (1), except that neurons now incorporate external inputs ***σ***_*t*_. They use these inputs to continuously sample the variable 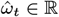 from the posterior *P* (*ω*_*t*_|***σ***_*t*_), and is read out through 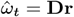. The details of *g*(**r, *σ***) depend on the choice of encoding kernel and noise statistics (Methods). In effect, this network takes velocity information encoded in Poisson spikes (Fig. 3B left) and continuously samples from the resulting posterior (Fig. 3B right, blue curve) through its spiking activity. While this signal seems to follow the true signal noisily (dashed curve), note that this variation reflects the actual shape of the uncertainty about the inferred variable (Supp. Fig. 3).

Notice that this implementation requires the network to continuously sample from a time-varying angular velocity posterior (Fig. 3B). Assuming that the dynamics of the variable encoded are slower than the neural dynamics, the network can draw multiple samples for the same angular velocity, allowing for a complete exploration of the posterior (Methods). In Supp. Fig. 3, we show how various statistics of the posterior distribution over angular velocities can be recovered from noisy Poisson inputs under the assumption that neural dynamics are 3 orders of magnitude faster than stimulus dynamics.

Overall, this implementation allows a spiking network to infer noisily encoded signals while capturing the full uncertainty over the posterior distribution.

### Inference, integration, and attractor dynamics within a single HD network

In this section, we will derive network equations for HD computations through sampling-based inference, which results in the integration of inferred angular velocity combined with attractor dynamics. Consider a population of Poisson neurons encoding angular velocity *ω*_*t*_ through spikes ***σ***_*t*_, and assume that the animal operates in darkness so that the only way to update the HD is through ***σ***_*t*_ inputs. Our derivation begins with the assumption that the network updates the current head direction estimate, 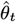, by drawing a sample from the distribution *P* (*θ*_*t*_|***σ***_*t*_, *θt*−1). This distribution can be computed with a statistical process called Bayesian filtering, which we approximate as follows (Methods): *P* (*θ*_*t*_|***σ***_*t*_, *θ*_*t*−1_) ∝ *P* (***σ***_*t*_|*θ*_*t*_, *θ*_*t*−1_)*P* (*θ*_*t*_|*θ*_*t*−1_) ≈ *P* (***σ***_*t*_|*ω*_*t*_)*P* (*θ*_*t*_). An intuitive interpretation of this process is a system that integrates angular velocity samples, 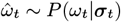 (Fig. 4A).

**Figure 4.**
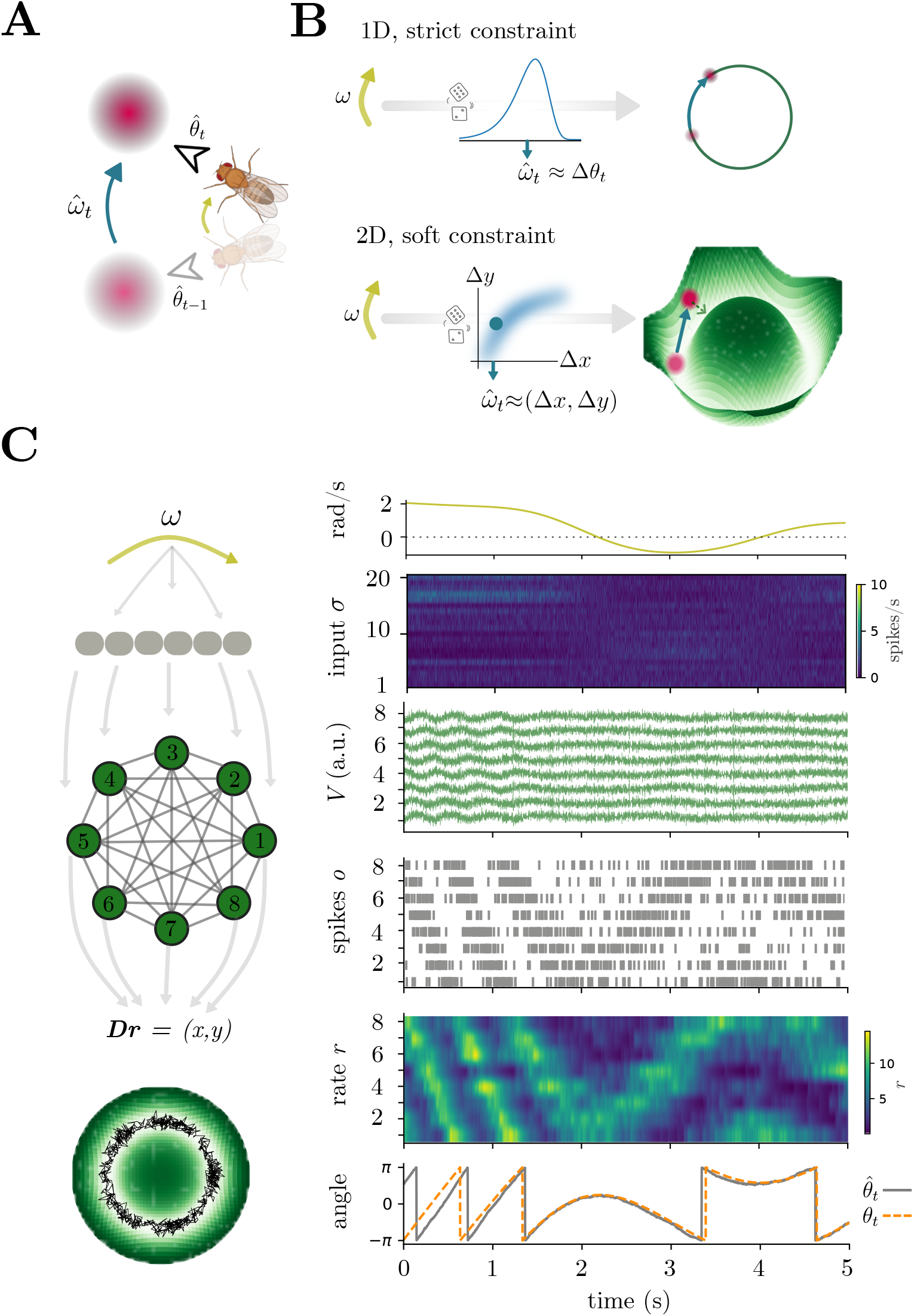
Combining sampling-based inference and attractor dynamics within a single network. **(A)** Updating the current HD estimate 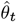 requires updating the previous estimate, 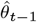, using 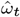, the angular velocity sample derived from input spikes ***σ***_*t*_. In statistical terms, this entails drawing a sample from the distribution 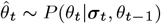. The distribution can be computed with a Bayesian filtering process *P* (*θ*_*t*_|***σ***_*t*_, *θ*_*t*−1_) ∝ *P* (***σ***_*t*_|*θ*_*t*_, *θ*_*t*−1_)*P* (*θ*_*t*_|*θ*_*t*−1_). We approximate this with the assumptions that 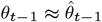 and *ω*_*t*_ ≈ *θ*_*t*_ − *θ*_*t*−1_. **(B)** Top: a possible neural representation consists of a 1 − *D* angle; in this case, the network is assumed to implement strict circular constraints, either through trigonometric functions or Fourier transforms. As described in the Methods, this implementation simply adds the current angular velocity sample 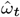 to the previous HD estimate. Bottom: another possibility is to consider a 2 − *D* representation of the angle. The usual transformation is *x* = cos(*θ*) and *y* = sin(*θ*); nonetheless, without a mechanism to pull the representation back towards *x*^2^ + *y*^2^ = *ρ*^2^, the network readout can go to zero or diverge. Hence, in our approximate Bayesian filter (Methods), we consider a specific prior to attract the network dynamics towards the circle. **(C)** Left: schematic of our implementation, with noisily encoded angular velocity, SCN with 8 neurons, and readout **Dr** (black curve) overlaid to the attractor prior (green ring). Right, top to bottom: angular velocity, noisy Poisson encoding ***σ***_*t*_, voltage of a SCN HD network with 8 neurons, spikes **o**, filtered spikes **r**, readout 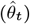 vs. real angle (*θ*_*t*_).

Head direction within the network can be represented in different ways, leading to different network mechanisms and dynamics (Fig. 4A-B, Methods). Here, we consider a network that outputs head direction estimates *θ*_*t*_ = (*x*_*t*_, *y*_*t*_) ≈ **Dr**_*t*_ in 2 − *D* coordinates. This representation prevents angular discontinuities and wraparound issues, can simplify downstream vector computations (e.g., navigation to a goal [77]), and could be used to represent other variables (e.g., forward speed). On the flip side, this representation requires a nonlinear mechanism to prevent the network output from drifting away from the circle (Supp. Fig. 5A). This can be done in different ways (Fig. 4A-B, Supp. Fig. 5B); here, we consider a prior that implements a soft circular constraint which, in the resulting dynamics, takes the form of an attractor (Fig. 4B, Supp. Fig. 5C). As shown formally in the Methods, we derive HD network dynamics of the form

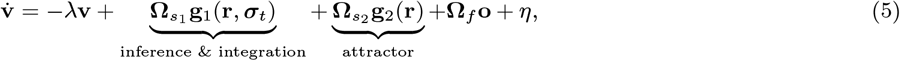

where 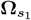 is a set of linear connections that infers information from external spikes ***σ***_*t*_, and 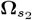 is a set of nonlinear interactions that attract the network activity to the circular manifold. Notice that, for clarity, we show the connectivities responsible for inference & integration and attractor dynamics separately in the equation; in practice, they are both part of the same slow connections responsible for sampling from a non-Gaussian probability distribution.

**Figure 5.**
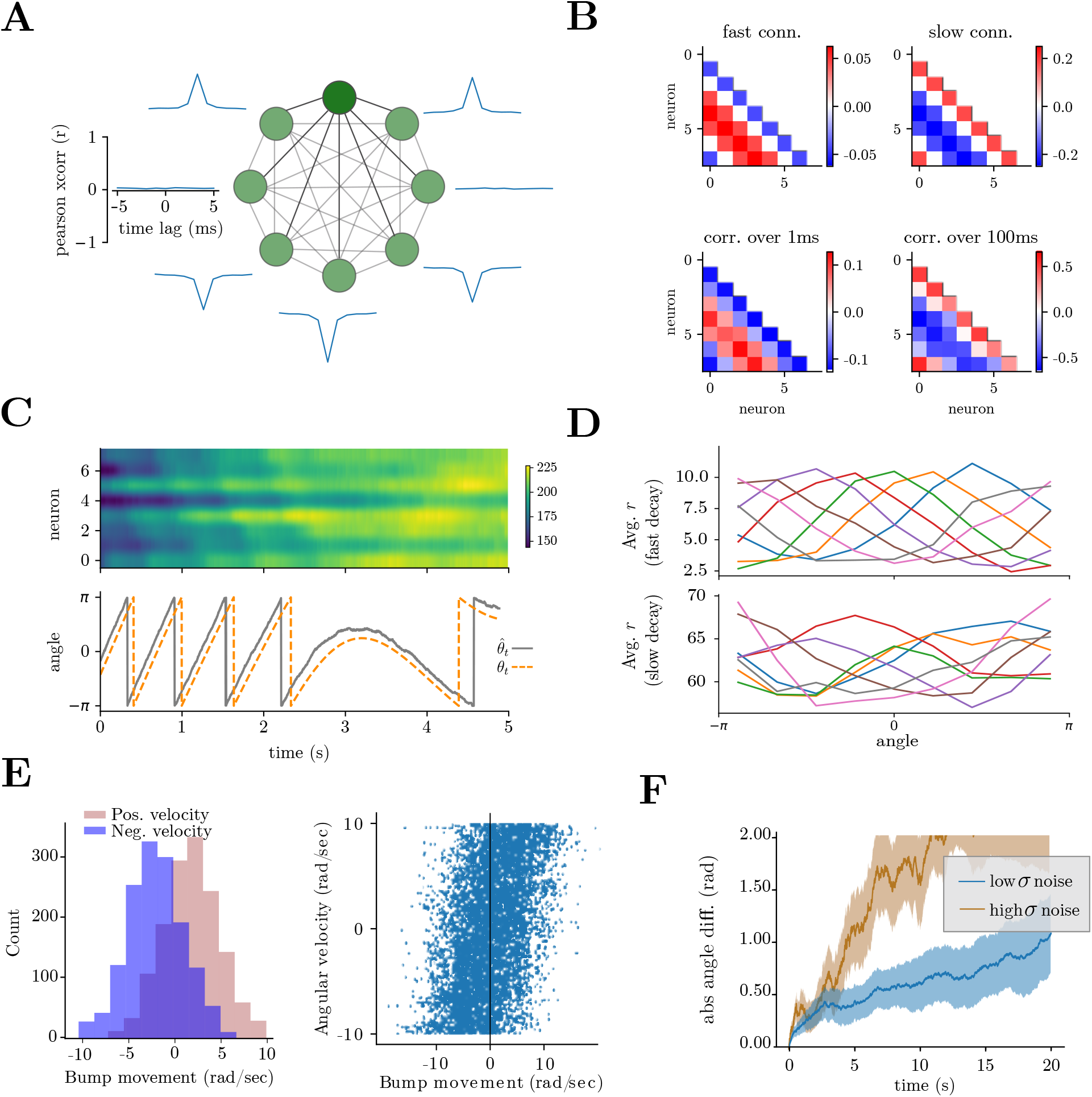
Experimentally testable predictions from SCNs performing HD computations through sampling. **(A)** Correlated stochastic fluctuations predict specific patterns of subthreshold voltage correlations among neurons with similar tuning. Top: example voltage trace for dark green neuron. Around the circle: subthreshold voltage cross-correlations between the green neuron and the others in the network. Notice that neurons with opposite tuning exhibit negative subthreshold correlations. **(B)** Top: fast connections show nearest neighbor inhibition and long-range excitation, while slow connections show a classical nearest neighbor excitation and long-range inhibition. Bottom: short-time vs. long-time correlations in neural activity, measured on spiking activity convolved with a Gaussian kernel of 1ms vs. 100ms. **(C)** Top: filtered spike trains **r** for a network with slow leak (*λ* = 0.5); Bottom: true HD orientation vs. network readout. **(D)** Tuning functions, measured as the average *r*_*i*_ given an HD angle *θ*, for networks of 8 neurons with fast leak (as in Fig. 4, *λ* = 10) vs. slow leak (*λ* = 0.5), as in panel C. Each line corresponds to one neuron. **(E)** Left: Distribution of bump velocity for positive and negative angular velocity (±4 rad/sec). The variability in bump movement velocity reflects the uncertainty in the inferred angular velocity. Right: Scatter plot of true angular velocity vs. bump velocity for a 50sec simulation. **(F)** Absolute angle difference during continuous clockwise turning at 5 rad/sec for low vs high input noise, averaged across 100 repetitions. Solid line: mean; shaded area: 1STD.

More specifically, these voltage equations implement the low-D dynamics in (*x*_*t*_, *y*_*t*_) defined by

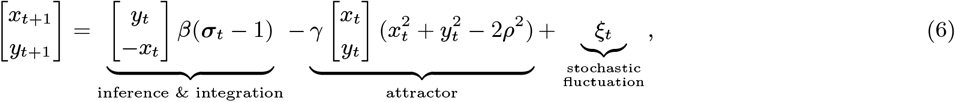

The first term, together with the stochastic fluctuations, approximates sampling-based inference from the posterior over angular velocity and integration towards the current HD estimate, while the second implements a soft constraint with strength *γ >* 0, which biases the dynamics towards the circle with radius *ρ* (Fig. 4B, Supp. Fig. 5).

Equation 5 constitutes the main result of our proposed SNN approach combining statistics with dynamics for handling uncertain angular velocity inputs. Starting from fundamental principles, we first derived networks capable of sampling from non-Gaussian distributions (Fig. 2 and Eq.1) to implement sampling-based inference (Fig. 3 and Eq.4). We then leveraged these results and extended them to derive a sampling-based ring-attractor network for inferring and updating the current HD representation given noisy velocity inputs (Fig. 4 and Eq.5).

Our derivation and implementation offer an alternative perspective on a computation necessary for effective navigation [3, 4]. Unlike previous approaches, our network explicitly accounts for the uncertainty of incoming signals, casting the problem as an inference rather than simply averaging the noise out. Such an approach opens avenues beyond HD estimation (as we discuss in the last results section) — for example, it enables downstream brain areas to directly asses the uncertainty in angular velocity inputs, which might be used for decision-making.

Finally, our network offers two key advantages. First, it is biologically plausible in terms of both connectivity and activity patterns, exhibits some well-known phenomena observed in real brains, and also generates novel testable predictions (as will be discussed in the final results section). Second, it is derived from well-known statistical sampling methods, making the model straightforward to adapt and extend, as demonstrated in the following subsection.

### Multimodal integration and visual reset

So far, we have considered only one external population input. In reality, the HD system receives a number of inputs [51, 22, 62]. This can be accommodated in our model by considering multiple populations of external inputs; consider *P* (***σ***^(1)^|*ω*), *P* (***σ***^(2)^|*ω*), …, each with their own encoding properties such as noise model, kernel, and nonlinearity (Methods). The angle update will follow

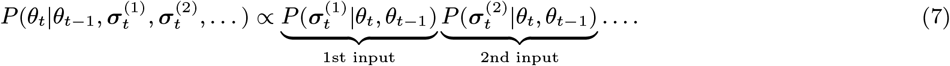

A relevant use case of multimodal integration is the implementation of a visual reset. As the network infers noisy information, the representation will drift due to noise [21], and requires periodic ‘reset’. This could, for example, be achieved by observing a visual landmark, allowing neural activity to realign with the true orientation. Such a visual reset is known to happen in the fruit fly, together with a plasticity mechanism that remapped visual stimuli onto the HD population by learning a specific angular relationship between the bump and the scene, called ‘visual remapping’ [14, 63]. We implemented a reset mechanism (Methods) through a second population of Poisson sensory neurons that gives a strong input to the network to move the bump in the right direction (Supp. Fig. 6A-B). We did not implement the plasticity mechanism, which could be implemented by taking inspiration from inhibitory Hebbian plasticity implemented in continuous RNNs [14]. While this implementation does not capture all biological complexities, it highlights the potential to extend the model to encompass a broader range of phenomena. Finally, in Supp. Fig. 1A, we demonstrate that our networks are not limited to sampling 1D variables but can sample from arbitrary distributions. This shows the potential for future extensions of our model to multimodal variable representations, inference, and integration.

### Experimentally testable predictions

Our HD sampling and attractor network (Eq. 5) reproduces several experimentally observed phenomena. Firstly, the network shows a localized “bump” of activity whose position correlates with the heading direction (Fig. 4C). The network maintains a good HD estimate for a period of time but slowly loses the true orientation as the various sources of noise accumulate (Supp. Fig. 4A, Fig. 5E), reminiscent of biological studies [21, 51]. Moreover, the average network activity is higher during periods of high angular velocity (Supp. Fig. 4B), as previously shown [21]. Finally, this HD system can be implemented in very small networks (Fig. 4, Supp. Fig. 4C) [20, 21].

Next, we use the model to generate experimentally testable hypotheses. Does the head directional system rely on sampling to infer angular velocity to be integrated? Does this process rely on an interplay between fast, slow, and nonlinear interactions, and are there correlated stochastic voltage fluctuations among nearby neurons? To answer these questions, we lay out hypotheses arising from our theoretical work and simulations and suggest concrete experiments and measures that could be studied in, but are not limited to, the *Drosophilla melanogaster* and the mouse HD systems.

We begin with low-level structural hypotheses:

#### 1. Voltage subthreshold fluctuations

To sample the full distributions, the neurons require specific subthreshold voltage fluctuations (Fig. 2). The voltages should exhibit stronger correlations among neighboring neurons. This can be tested by computing cross-correlations between the voltages of nearby and distant cells (Fig. 5A) and could be achieved experimentally, for example, in a fly navigating an artificial environment paired with voltage imaging of the central complex neurons [21, 78, 79], or head-fixed mice navigating in an artificial environment [79]. Moreover, given the absence of sensory stimuli during the larva/pupa stages of fruit fly development, one could study the dimensionality of neural activity, which will be driven by other areas’ inputs and other noisy sources, which, summed, could be seen as structured voltage noise. This prediction derives from mapping the Langevin sampling dynamics to a neural network (Fig. 2B right).

#### 2. Fast vs. slow interactions

The networks presented in Fig. 4 have two main sets of connections. The fast connections, which operate on a timescale of a few milliseconds, show nearest-neighbor inhibition and long-range excitation. On the other hand, slow connections, which operate on the timescale of tens of milliseconds, show a classical nearest neighbor excitation and long-range inhibition, similar to what was used in previous studies [49], inspired by true biological connectivity [62] (Fig. 5B, top). This is observable in the short-time versus long-time correlations in neural activity (Fig. 5B, bottom), an analysis that could be carried out using ultra-fast calcium recording [80] or e-phys recordings in the central complex of fruit fly, or with detailed knowledge of the timescales of interactions in a connectome [62]. Nonlinear interactions will act over slow timescales and constrain the activity to the desired manifold, and the manifold dynamics can be made arbitrarily slower at the cost of softening the circular constraints *γ* in Eq. (6). This is biologically relevant, as these slower nonlinear interactions might be implemented through widespread peptidergic neuromodulation [81], and could be studied in a peptidergic connectome [82]. This prediction comes from using SCN theory to implement the gradient dynamics required for Langevin sampling (Fig. 2B left).

Next, we look at network-level hypotheses:

#### 3. Angular integration without bump selectivity

Some values of voltage leak term *λ* lead to a loss of the bump in neural activity, whereas the linear read-out 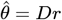 remains constrained to a circle (Fig. 5C). This is reminiscent of HD systems in vertebrates, where the neural activity does not show an obvious bump of neural activity, although the low-D manifold representation clearly falls on a circle [3]. Moreover, the resulting circular tuning functions are not necessarily equal across neurons (Fig. 5D), similar to what is observed in rodents, and that violates the assumptions of classical ring-attractor models [83, 84]. Experimentally, one could observe that the absence of a bump of activity due to the slow voltage leak increases tuning function heterogeneity but does not hinder the readout quality. This could be achieved, in principle, through genetic or pharmacological interventions to manipulate specific ion channels in fly models with slow vs. fast voltage leaks of central complex neurons. According to theory, the bump should disappear without hindering HD readout precision. This prediction derives from the encoding properties of SCNs.

#### 4. Bump movement statistics

In our model, the head direction is updated by sampling from the posterior over angular velocities; this implies that changing informativeness of the inputs will change the certainty of updates (i.e., the bump will exhibit bigger vs smaller “jumps”) (Fig. 5E). This can be tested in various ways: for example, one can manipulate the reliability of GLNO input neurons or provide conflicting visual vs. locomotive stimuli [1], with simultaneous recordings from central complex neurons. Such analyses are not currently feasible due to slow calcium sensors and altered behavioral/proprioceptive dynamics due to head-fixation and fly-on-a-ball setup [49]. In the future, faster sensors and free navigation might provide an avenue for implementing such analyses. This prediction comes from the use of probabilistic sampling to infer head direction.

#### 5. Skewed Poisson posterior

Given that the Poisson posterior is skewed, and the skewness depends both on angular velocity and input quality, it is possible to observe the same skewness in the statistics of bump movement. In particular, comparing the instantaneous bump position finite difference vs. fly angular velocity, it is possible in our model to visualize the skewness of the distribution for positive and negative angular velocities (Fig. 5E). As in the previous point, these measures are unreliable in current experimental settings and will necessitate faster sensors for future exploration. This prediction is a result of assuming the velocities are encoded through Poisson spiking and then sampled from the posterior distribution by the HD network.

Finally, we propose functional and behavioral hypotheses on and around the question: “Does explicit knowledge of input uncertainty help, functionally and behaviorally?”. These predictions all link to the assumption of a network which explicitly uses probabilistic sampling to estimate and infer information.

#### 6. Fixed vs flexible mapping of posterior to velocity estimation

The simplest option is to have a hardwired circuit that automatically converts inputs into velocity estimates without stochastic sampling. Nonetheless, the posterior might change significantly during behavior, either due to noise or differing input statistics, suggesting a need for a flexible, probabilistic adjustment mechanism. These two possibilities could be disentangled experimentally by manipulating input firing properties or input statistics and then measuring bump movement statistics as a proxy for studying posterior distributions. In particular, a fixed mapping will maintain a fixed posterior shape, whereas a flexible one should be able to adapt.

#### 7. Unreliable inputs and angular velocity estimation

The noise of the inputs determines how quickly the network will lose its true orientation (Fig. 5F). If inputs from certain neurons (e.g., GLNO neurons in the fly brain) are unreliable, P-EN neurons may rely on other cues for angular velocity estimation. For example, recent work by [1] shows that unreliable GLNO inputs force P-EN neurons to use other cues to estimate angular velocity. This situation could illustrate how the HD system uses estimates of input uncertainty—such as higher moments of the angular velocity posterior—to switch its reliance on different cue modalities. One possible experiment would be to measure the bump movement sizes and determine whether they correlate with GLNO input uncertainty only up to a particular threshold.

#### 8. Behavioral relevance of input uncertainty

Does the explicit knowledge of uncertainty in sensory inputs affect behavior? This question could be addressed by manipulating input conditions or by observing behavior under conflicting proprioceptive and visual cues. For instance, if specific angular velocity encoding neurons are disrupted or manipulated, does the subject exhibit less turning behavior or rely more on non-HD cues? Additionally, understanding whether the explicit tracking of uncertainty is advantageous in changing environments—such as variations in lighting or input quality, damage to neurons, or altered sensory modalities—could provide valuable insights into adaptive behavior.

## Discussion

The brain is hypothesized to use sampling-based methods to deal with uncertain inputs, but important open questions remain about how local interactions and neural variability come together to perform such network-level probabilistic computations in non-sensory brain areas. We described the neural and structural requirements for spiking HD neural networks to combine sampling-based inference and attractor-based integration. To this end, we described the relationship between nonlinear synaptic interactions and sampling from exponential family distributions, sampling-based inference from noisy stimulus encoding, and sampling-based integration of noisy angular velocity inputs. We concluded with biological hypotheses and experimental predictions derived from this framework.

### HD neural computations as sampling-based inference

Classical models of HD ring attractor networks [45, 48, 47] often rely on strict assumptions about the input signals, and generally do not cast decoding the inputs as an inference problem. How should a spiking neural network deal with generic, noisy inputs? A previous study by [53] proposed using a circular Kalman filter to implement a version of Bayesian ring attractor. In that implementation, the network keeps track of mean and circular variance, and the bump modifies its amplitude in response to confirmatory vs. conflicting evidence. In the invertebrate brain, the bump does not seem to change in size during darkness [21, 22, 49], suggesting that the head direction system does not directly represent uncertainty by widening the activation bump. Nonetheless, uncertainty should not be discarded so readily. Empirical evidence suggests that angular velocity inputs are noisy [22, 2, 1], and the head directional system must perform computation on this input to infer the most likely angular speed. In our work, we assume that the HD system can deal with the noise by performing sampling-based inference. The network we derive combines aspects of statistical inference and classical attractor network dynamics. Its behavior can be understood as integrating samples from the posterior distribution over angular velocities while maintaining a soft circular constraint. Under the assumption that neural dynamics are faster than behavioral dynamics, this leads to the integration of the posterior mean. Moreover, this approach guarantees the possibility to compute arbitrary statistics on the inputs. These capabilities could be useful in cases where the posterior changes over time or across conditions, and future work will probe for flexibility and robustness through plasticity rules adapting this sampling to different inputs. In our study, the bump does not significantly change its shape during periods of increased uncertainty. Rather, it “wiggles” around the true head direction due to the Poisson inputs and the randomness driving sampling. Moreover, [53] predicted that in the absence of visual inputs, the network would progressively lose direction selectivity. In our study, although the true angular direction becomes imprecise over time, the bump selectivity does not decrease. Other aspects of our simulations are consistent with previous reports. For example, the bump size does not significantly change throughout the experiment, the average activity is higher during period of high angular velocity, and the true orientation is progressively lost in darkness [21, 22]. Moreover, we need just a handful of neurons for the precise integration of angular velocity inputs, as is known to happen in several insect brains [85], and recently demonstrated in a study on rate network attractors [49]. Finally, the slow connectivity of our network is consistent with previously proposed ring attractors, as in nearby excitation and long-range inhibition [48]. Although we do not claim that our model is a faithful reproduction of the head directional system, we hope for it to be an alternative viewpoint and inspiration for future experiments. To that extent, we provide extensive predictions related to the connectivity and neural activity correlations over different timescales, subthreshold inputs, and other testable measurements.

### Alternative spiking ring attractor networks

We briefly review other models of spiking neural networks implementing ring attractors in the context of existing literature.

[86] proposed a general framework for biologically plausible simulations of attractor networks, including the HD system, based on the Neural Engineering Framework [87]. The paper’s main idea is to implement control-theoretic approaches in networks of spiking neurons; for the ring attractor, a nonlinear control model is built using a vestibular control signal to move the current head direction estimate. An interesting contrast to our model is that they explicitly avoid the use of multiplicative interactions at the neural level, while in our model, this is the enabling factor. Moreover, they make use of an explicit notion of tuning function for each neuron, and introduce nonlinearities at the level of encoding external variables, while preserving the notion of linear decodability. A limitation of their approach is that the decoding weights need to be learned through least-squared methods, and it is not clear that this HD system can be implemented with a handful of neurons only, both of which can be easily solved with our approach.

[88] built a detailed spiking ring attractor model, striving to computationally mimic the *Drosophila melanogaster* head direction system [21, 89]. The protocerebral bridge (informing of angular velocity) and ellipsoid body (ring attractor, maintaining bump of neural activity) are modeled explicitly, as well as the neurons projecting between the two networks, using leaky-integrate-and-fire (LIF) dynamics. Most single neuron parameters correspond to experimentally observed values, while neural connectivity strengths were set through parameter sweeps. Their results successfully replicate several findings from [21] and interestingly suggest that spiking neurons are required to recapitulate other aspects of fruit fly experimental recordings. In the absence of spike inputs and external cues, bump movements were completely absent. In contrast, the bump would still move in our model (and also in the presence of reset mechanisms, the bump would still “wiggle”) because of the subthreshold voltage stochastic fluctuation and internal spiking activity implementing sampling.

Finally, work by [52] built a network model for the mammalian lateral mammillary nucleus and dorsal tegmental nucleus (LMN-DTN), consisting of three populations of noisy and spiking neurons, coupled by biophysically realistic synapses. The angular velocity updates for the head direction were fed into the network as uncorrelated Poisson spike trains, specifically to neurons in the inhibitory populations - consistent with experimental findings [2]. An interesting finding with respect to the bump of neural activity is that the external excitation, together with recurrent cross-inhibition was sufficient to produce the persistent activity, without the need for recurrent excitation. In contrast, our model requires the slow connections to have nearest neighbors excitation to preserve the current HD estimate.

### Structural and neural requirements

In our networks, we formalize sampling as Langevin dynamics; this method combines the gradient of the log-probability with a source of stochasticity to explore the probability landscape. What are the neural requirements at various levels, and how biologically feasible is it? The gradient of the log-probability is, in most non-Gaussian settings, a nonlinear function, which requires the use of nonlinear interactions to be implemented or approximated. In our work, we formalized multiplicative synapses as the ideal set of interactions for exponential distributions with polynomial moments, which can be used to approximate generic distributions to arbitrary precision[90]. The feasibility of this type of interaction has been discussed at length in [56]. As a summary: 1) experimental circuit reconstruction studies demonstrated the existence of several distinct synaptic connections among cell pairs [91, 92]; 2) examples of effectively multiplicative computations have been characterized [93, 94, 95, 96]; 3) experimental and theoretical work has long shown the computational advantages of nonlinear synaptic and dendritic interactions in single neurons [97, 98]; 4) mechanisms such as dendritic calcium or NMDA spikes, synaptic clustering, and shunting inhibition [99, 100, 101, 102, 103, 104] could all contribute to implementing effective multiplicative interactions. Recently, the biophysical details for implementing multiplication-like nonlinearity have been described in a class of *Drosophilla melanogaster* ON motion vision neurons, which arises from the coincidence of cholinergic excitation and release from glutamatergic inhibition [105]. An alternative formalism to approximate multiplications is the use of nonlinear dendrites, which have been shown to act as universal function approximators [106], also in spiking neural networks [68, 71]. Moreover, it was shown previously that it is possible to employ supplemental networks in order to avoid high-order interactions [56] or to simplify certain computations taking advantage of the fixed points of recurrent networks [107, 34]. Finally, all connections will depend on the choice of sparseness of the decoding matrix; the connectivity density of *g*−th order interactions will behave as ∝ (*pN*)^*g*^ [56].

Next, the assumption of Langevin sampling requires the existence of correlated voltages/stochastic subthreshold fluctuation between the neurons. Neural variability at various levels has been observed in many studies. The inherent variability of neural responses to identical stimuli has been observed for decades [108, 109, 110, 111]. This, together with the presence of various types of noise both at the sensory, cellular, synaptic, and population levels [11], have been interpreted as support for probabilistic neural representations [30, 24] and for the idea that these sources of variability facilitate the exploration of posterior probability distributions over stimuli [35, 5, 7, 39]. In particular, and most relevant for our implementations, is the need for correlated stochastic fluctuation, which are extremely important in our network as they are the principal driver of sampling. This requirement gives rise to consistent ongoing subthreshold synchronization without the need for oscillations and stronger correlation in pairs of neurons with similar receptive fields [72, 73, 74, 75].

#### Other approaches to sampling in spiking neural networks

Various approaches have been proposed to explore sampling methods within Spiking Neural Networks (SNNs). Among the prominent techniques, Markov Chain Monte Carlo (MCMC) methods utilize stochastic neurons to approximate samples from target random variables, offering a robust framework for probabilistic inference [35, 36, 37, 38]. In our case, single neurons don’t necessarily represent sampling proposals, and aren’t required to work with discrete probability distributions. More recently, a fractional diffusion neural model has been proposed that harnesses Levy motions to effectively sample from heavy-tailed, non-Gaussian distributions [40]. Although biophysically realistic, this model relies on many thousands of excitatory and inhibitory neurons embedded in a 2*D* feature space, and it is not clear how the complexity of the neural circuit model and the sampling dynamics would be affected by increasing the dimensionality of the problem. Another recent approach that used different assumptions involves the use of linear recurrent networks of excitatory neurons governed by Poisson statistics [39]; these networks don’t need external correlated stochastic fluctuation in the form of a Wiener process, but instead rely on the intrinsic Poissonian variability of the inputs to drive the random sampling from the stimulus posterior [39]. Finally, as in our work, previous implementations of Langevin samplers relied on SCNs [41, 42]. These networks can be fully derived and embed various types of deterministic and stochastic dynamics directly into their computational architecture [54, 55], including Langevin Dynamics [41]. Importantly, [42] have bridged the gap between generic MCMC-based spiking neurons and SCNs, revealing a deeper connection between these two sampling SNN frameworks.

### Closing remarks

We have explored how sampling-based computations and attractor dynamics can be implemented within spiking neural networks in the special case of the HD system. This work supports the hypothesis that sampling-based methods are a viable and powerful framework for neural computation in non-sensory brain areas, offering hypotheses for experimental and computational neuroscience. Future research will further explore these concepts, particularly in more realistic and higher-dimensional settings, and integrate findings with experimental data to enhance our understanding of stochastic computations in the brain.

## Methods

### Notational convention

First we introduce the symbols used throughout this work, and specify dimensionalities or values used

**Table 1.**
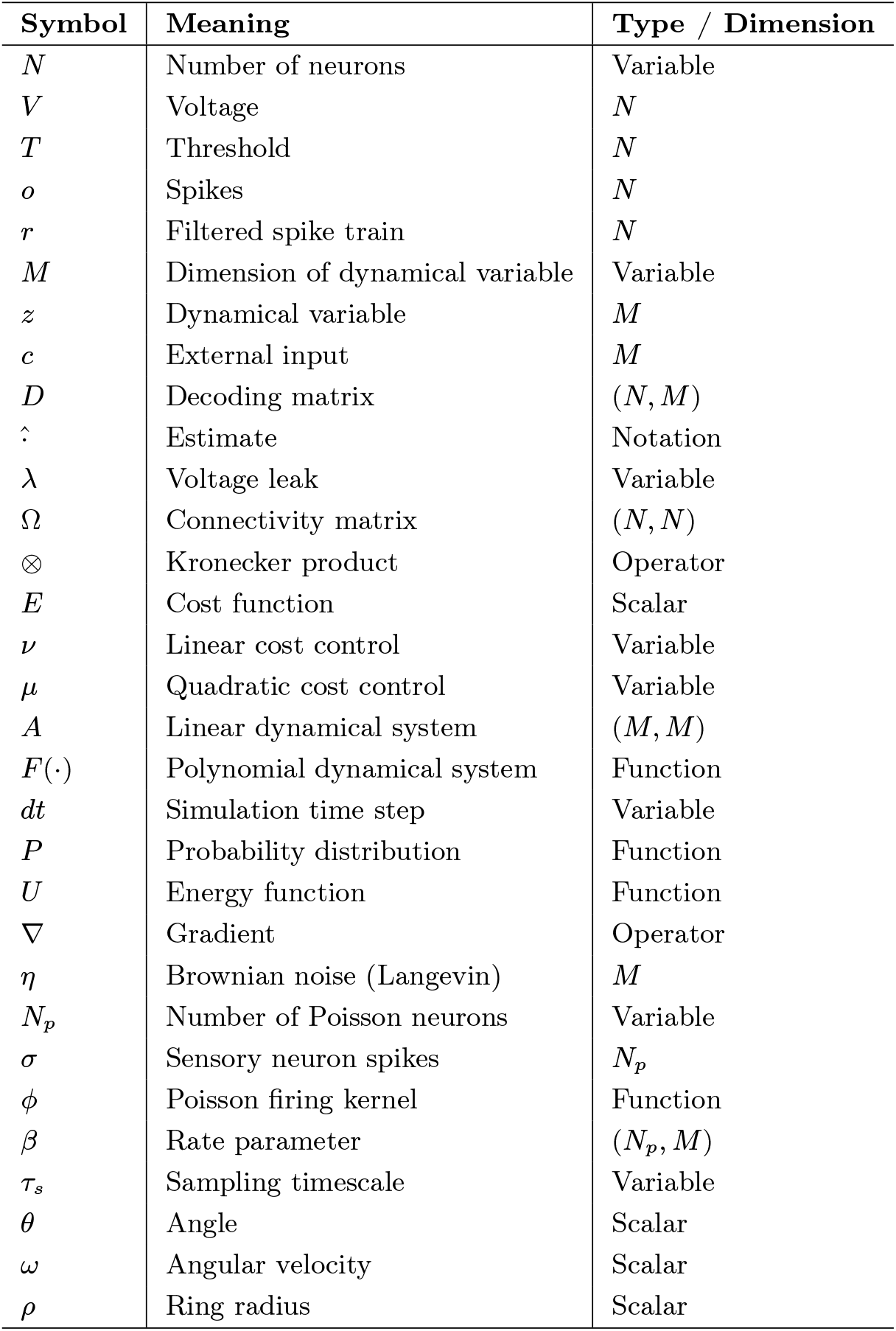
Notational Conventions.

### Spike Coding Networks

We start by introducing the model by [54], upon which we will expand in the coming sections.

The goal is deriving the equations of a network that follows the temporal evolution of a dynamical signal **z** ∈ ℝ ^*M*^,

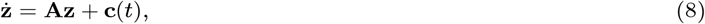

where **A** ∈ ℝ^*M*×*M*^ is the state-transition matrix and **c** ∈ ℝ^*M*^ is the external input. The network readout, 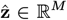, is an estimate of the signal **z**,

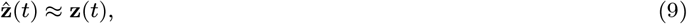

and is updated dynamically using the spiking of the *N* neurons, **o**(*t*):

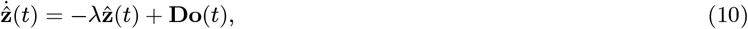

where *λ* is the leak term, and **D** is an inherent and fixed decoding weight matrix of size *M* × *N*. For simplicity, we will consider the vector **r** of filtered spike trains

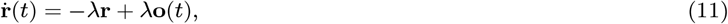

such that the stimulus estimate

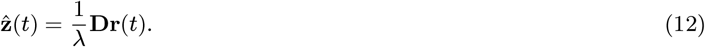

To minimize the difference between the real signal **z** and the readout 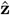, the cost function *E*(*t*) is minimized

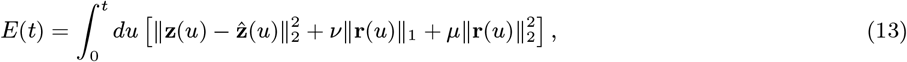

where the first term minimizes the difference between signal and estimate, the second term forces to minimize the amount of spikes, and the third term distributes spikes across neurons, leading to efficient and balanced spiking, respectively.

Consequently, the network dynamics will perform greedy minimization of *E*(*t*), such that they only spike if it leads to a decrease in *E*(*t*). The spiking rule for neurons will be

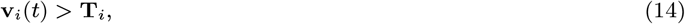

where

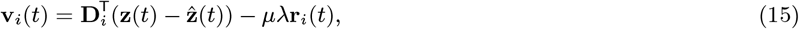

and

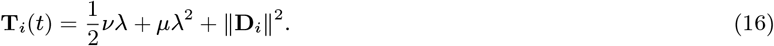

With *ν, µ* = 0,

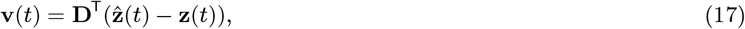

and

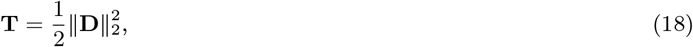

Taking the time-derivative of **v**, rearranging, and using the approximation 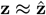, we obtain [54, 41, 58]

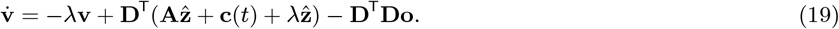

### Multiplicative interactions in SCNs

If the dynamics of **z** are described by a linear system,

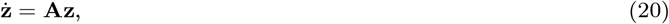

we can re-write eq. Eq. (19) as

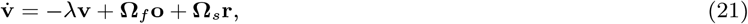

where

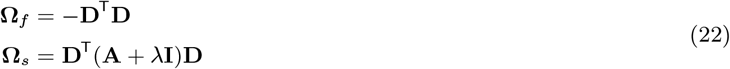

are the fast and slow recurrent connectivity matrices, respectively. In the case of non-linear systems of the form **ż** = *F* (**z**), the resulting model has formal network dynamics given by

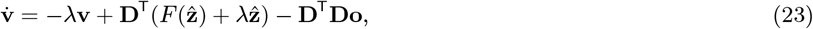

where *F* (·) is a non-linear function. One approach is to approximate the function *F* by using a polynomial expansion up to order *g*:

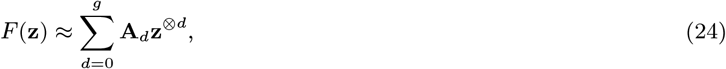

where ⊗ denotes the Kronecker product (see below). This derivation quantifies the minimum set of multiplicative interactions [99, 95, 104, 96, 94, 93, 105] necessary for implementing a polynomial dynamical system in a SCN, which we call multiplicative SCN (mSCN) [56]:

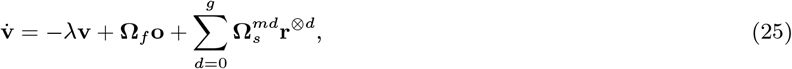

where

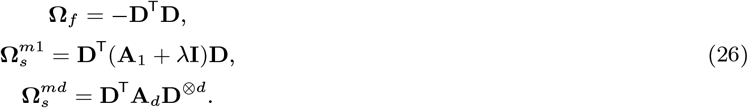

All these interactions could be carried out in the dendritic tree of a neuron [97, 112].

Next, to simplify notation and intuit computational implementation, we rewrite the summation to a vectorized form through concatenation. First we define the term **Ω**_*s*_ that encapsulates all the interaction matrices 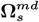 as

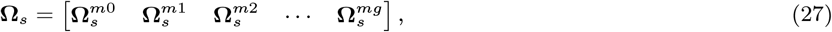

where each 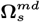 corresponds to the individual interaction matrices defined in Eq. (26). Furthermore, we introduce the term *g*(**r**) ∈ ℝ^*L*^ as a function of **r** that, in our case of multiplicative interactions, contains all the powers of **r**, such that

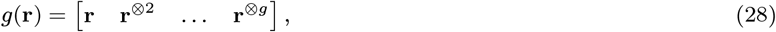

This term represents all the possible multiplicative interactions in **r**, from **r**^⊗1^ to **r**^⊗*g*^, and *L* is of length 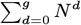. This allows the expression of the sum in the compact form,

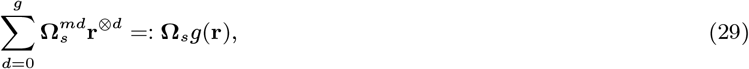

which leads to the resulting expression for the voltage equations

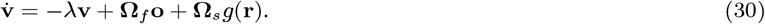

Note that *g*(**r**) could consist of other types of non-linear interactions as well, depending on the method at hand and the capabilities of the neurons in the network [97].

#### Kronecker formalism

Multiplicative interactions in the SCN framework depend on the use of the Kronecker product. The ⊗ symbol represents the Kronecker product, of which the operation between two matrices is given by

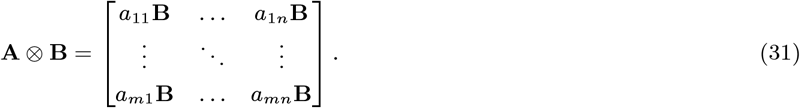

An interesting property of the Kronecker product is the mixed-product property, given by

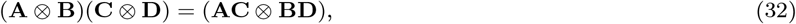

which is allowed in the case that the matrix products **AC** and **BD** exist. As in [56], we will often use the notation **D**^⊗*d*^ to denote the *d*-fold Kronecker product of **D** with itself; i.e. if *d* = 3, then **D**^⊗3^ = **D** ⊗ **D** ⊗ **D**. Summarized biological support for multiplicative interactions can be found in the Discussion and in-depth elaboration is to be found in [56]

### Sampling using SCNs

The classical SCN formulation [54, 58] is given by

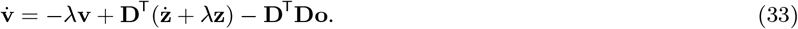

On the other hand, the general form of naïve Langevin Dynamics is given by

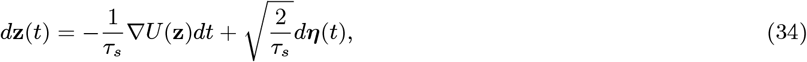

where *U*(**z**) is the log-probability density function (PDF), *τ*_*s*_ is the timescale of the sampling performed by the network, and ***η*** ∼ N (0, **I**) is Brownian noise. Specifically, we consider PDFs belonging to the exponential family,

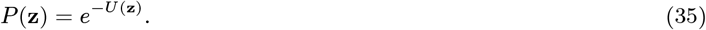

The Langevin Dynamics Eq. (34) can be formally plugged into the membrane voltage equation Eq. (33), as in [41, 42], to obtain

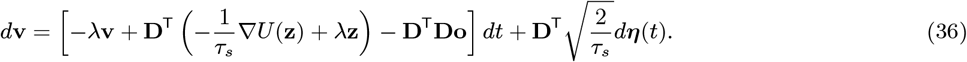

Lastly we use the assumption that 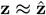,

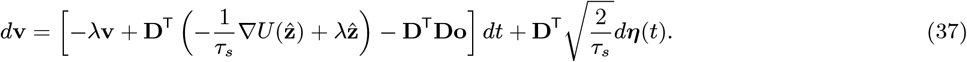

For the Gaussian distribution, the gradient of the energy function will be linear in **z** and consequently also linear in **r** [41, 42]. However, as is, the current formulation does not accommodate more complex distributions, as the energy function will contain non-linear interactions between **r**. Note that in the main text, for clarity, we used ***η*** for the voltage noise Eq. (5), whereas ***ξ*** was used for the stochastic fluctuations pertaining to the Langevin Dynamics Eq. (6); practically, these were related through ***η*** = **D**^**⊤**^***ξ***, where ***ξ*** is projected from the dynamical variable dimension *M* to neural dimension *N* through **D**. In all of the Methods, ***η*** takes the place of *ξ*, and the projection is explicitly written, as seen above.

### Sampling polynomial-moment exponential family distributions

Consider *P* (**z**) ∝ *e*^−*U*(**z**)^ for **z** ∈ ℝ^*M*^ ; if *U* (**z**) is a polynomial with maximum degree *g* + 1, ∇_**z**_*U* has maximum degree *g* and can always be written in the form

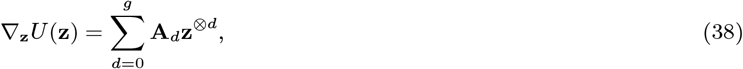

where 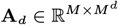 is the matrix of coefficients for the degree-*d* terms in ∇*U*, and we define **z**^⊗*d*^ as above. Then we can rewrite Eq. (37) as

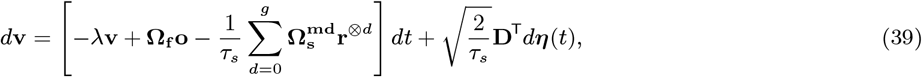

where

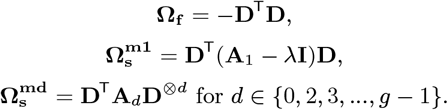

Notice that we must ensure that the polynomial *U* (**z**) is well behaved, i.e. that ∫*e*^*U*(**z**)^ *d***z** *<* ∞. This is true, for example, in 1 − *D* for polynomials *U* (**z**) with a maximum degree that is even, and when the highest order term has a negative coefficient (e.g. *U* (*z*) = −*z*^2^).

Using the same notation introduced in Eq. (29), we can rewrite Eq. (39) to the equation found in the main text Eq. (1):

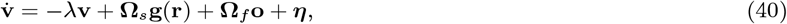

### Sampling-based inference of stimuli through Poisson sensory neurons

A latent variable is encoded by noisy spiking neurons, and the spiking is then used by downstream regions for further computation. One such computation is the inference of the latent variable, which will be inherently uncertain due to the encoding noise. One way to deal with this uncertainty is to sample from the posterior distribution over latent variables, a process called sampling-based inference [41, 26].

Starting with a single Poisson neuron with firing rate kernel *ϕ* dependent on the incoming stimulus **z**:

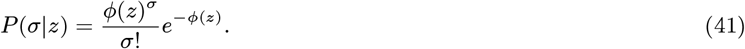

The posterior distribution over latent variables, assuming a flat prior, can be obtained with Bayes’ rule simply as

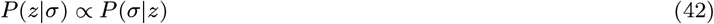

Here, we want to sample from *P* (*z*|*σ*) using Langevin Dynamics, which is, formally:

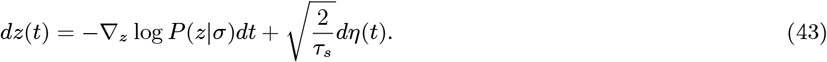

Extending to multiple neurons and high−*D* variables **z**, we can write

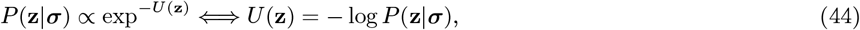

where

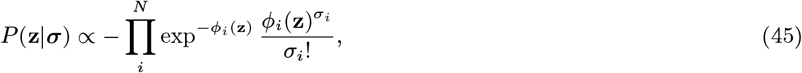

such that

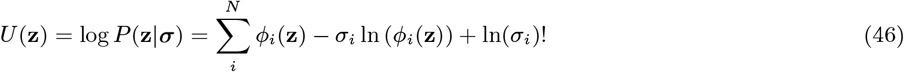

Consequently,

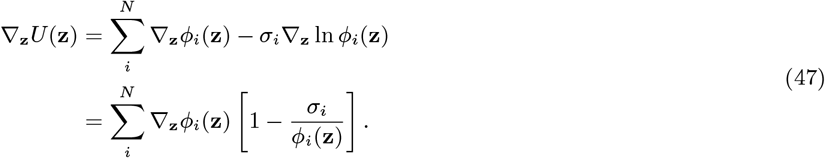

Once a kernel is chosen, Eq. (47) can be plugged into Eq. (37) to obtain a SCN that performs sampling from the posterior distribution over latent variables **z**, which is read out through 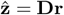:

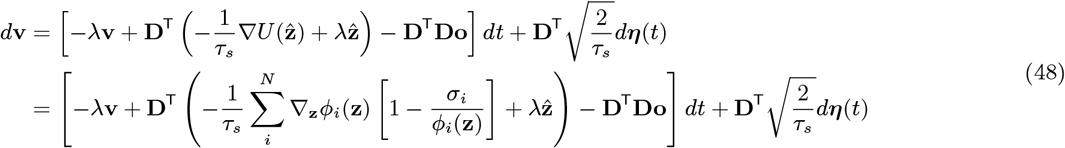

In the following, we will see the different forms of this equation for different choices of firing rate kernel *ϕ*, but in principle, any positive function with continuous derivative will work.

#### Exponential Kernel

Choosing an exponential firing rate kernel,

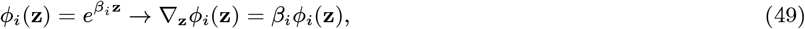

such that

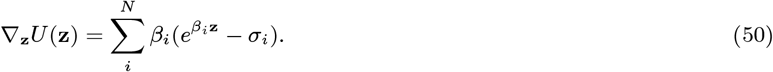

we find Langevin Dynamics on **z** evolve according to

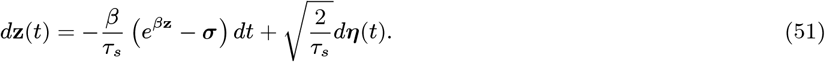

We can now plug this into the membrane voltage equation Eq. (48), discerning between the spiking of the Poisson neurons ***σ*** and the spiking of the mSCN **o**, and using the fact that 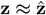,

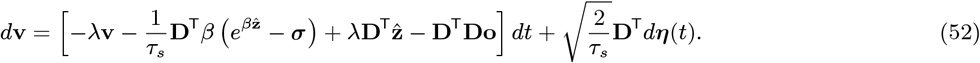

Using 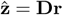,

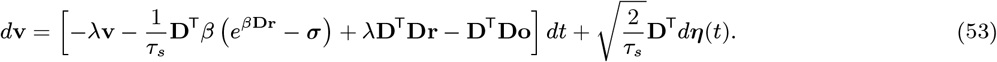

The exponential term can be approximated by its truncated Taylor series and rewritten with multiplicative synapses as in the main text Eq. (1).

#### Linear Kernel

For a linear firing rate kernel with base firing rate *r*_0_,

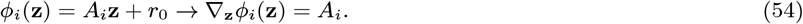

Plugged into Eq. (47),

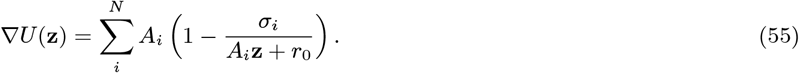

Using the simple approximation 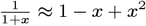,

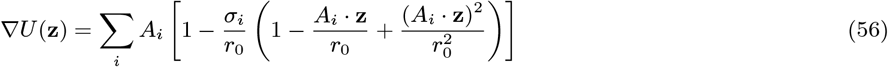

If we use 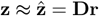, then we can rewrite (*A*_*i*_ · **z**)^2^ as

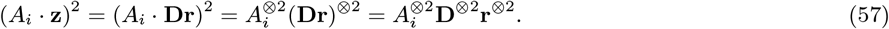

Plugging this into the equation for ∇*U*,

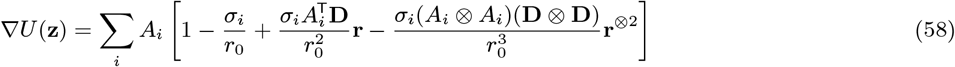

Writing in vectorized form and plugging into Eq. (48), one finds

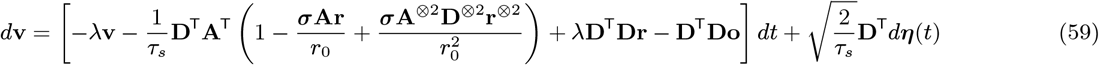

This derivation can be extended to arbitrary precision by including more Taylor expansion terms, up to *K*:

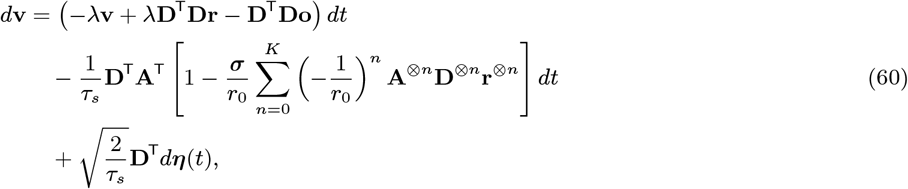

#### Logistic kernel

For a logistic firing rate kernel

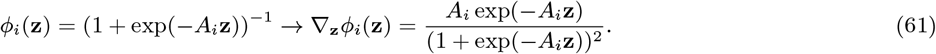

Plugged into Eq. (47),

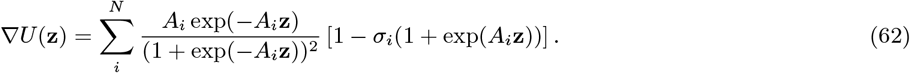

Let’s consider for simplicity the case where **z** is 1 − *D*; similar approximations can be achieved in higher dimensions. The term ∇_*z*_ *ϕ*_*i*_(*z*) is well approximated, within a neighborhood of *z* ≈ 0, by a parabola:

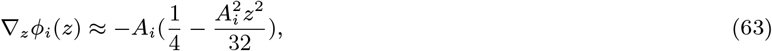

therefore we consider

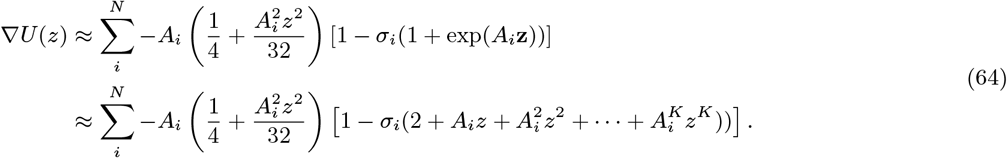

This is a polynomial of order *K* + 2, and can be easily implemented into Eq. (48).

#### Different noise models

Other discrete probability distributions that can be chosen as noise models and implemented in Eq. (48) upon choosing appropriate approximations and/or synaptic nonlinearities:

##### Bernoulli

Suppose we are dealing with independent binary neurons *σ*_*i*_ with firing probability sigmoid *f*_*i*_(**z**) ∈ [0, 1]:

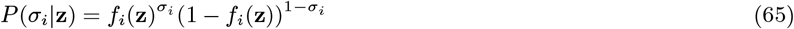

Suppose that 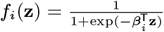. Taking the gradient of log *f* (**z**) we have the simple relationship ∇_**z**_ log *f* (**z**) = *β*(1 − *f* (**z**)), hence

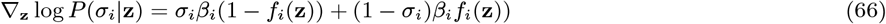

Considering now a population ***σ*** = {*σ*_1_, …, *σ*_*k*_} where *P* (***σ***|**z**) = ∏_*i*_ *P* (*σ*_*i*_|**z**),

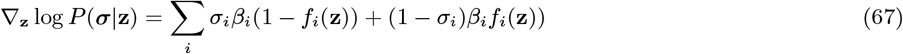

##### Binomial

If each neuron is binomial, as in *σ*_*i*_ ∈ {0, 1, …, *n*} and

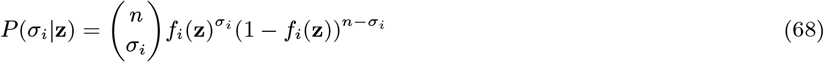

there follow similar derivations as above, and

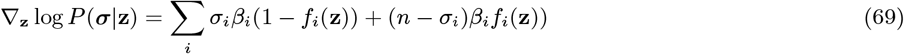

##### Geometric

if *σ*_*i*_ ∈ {0, 1, …}, where 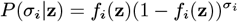, then

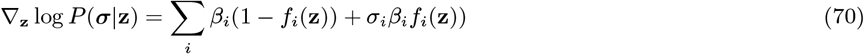

Separation of timescales, and temporal integration to estimate moments

To deal with dynamically encoded stimuli, we assume that the neural dynamics are at least one order of magnitude faster than the stimulus dynamics; this could be justified by observing that animals interact with their surroundings with actions often spanning hundreds of milliseconds, while the average membrane timescales in the cortex are on the order of tens of milliseconds; or by observing the millisecond precision of stimulus-related spikes in the visual system of multiple organisms for natural stimuli with much slower timescales [113, 114]. This allows for the exploration of the posterior distribution over dynamic stimuli online, through a moving average or a similar filter.

### Inference of Heading Direction through sampling-based integration of angular speed

We want our network to perform inference on the current head direction given the previous estimate and the angular velocity provided by the external neural population. We assume that the head direction estimate of our network will be given by

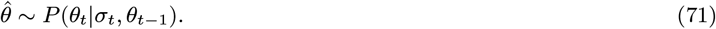

This estimation can be decomposed using Bayes’ rule:

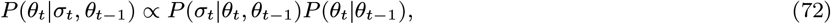

We consider the case where *σ*_*t*_ depends only on the change in angle, Δ*θ* ∝ *ω*_*t*_, and assume that

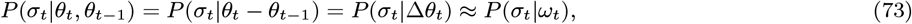

such that Eq. (72) becomes

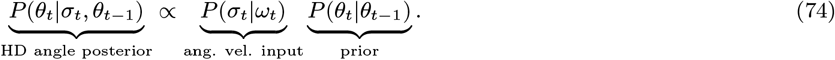

The term *P* (*σ*_*t*_|*ω*_*t*_) will inform the HD system about the angular velocity *ω*_*t*_ through Poisson spikes *σ*_*t*_, while the term *P* (*θ*_*t*_|*θ*_*t*−1_ will implement a prior, which can take different forms, as described below. Finally, Langevin sampling in *θ*_*t*_, written in a discretized form amenable to being implemented in SCNs, and assuming *ω*_*t*_ ≈ *θ*_*t*_ − *θ*_*t*−1_ ≪ 1, *P* (*σ*_*t*_|*ω*_*t*_) ∼ *P oiss*(exp(*βω*_*t*_)) and a flat prior:

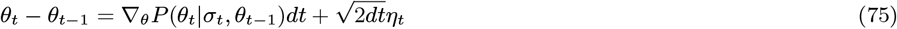

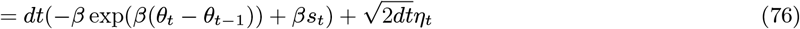

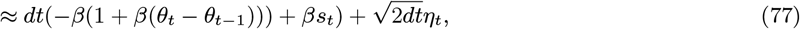

where *η*_*t*_ ∼ *N* (0, 1). Denote *α* = 1 + *β*^⊤^*βdt*, then ^1^

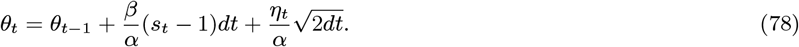

This equation could be implemented in a SCN, e.g., Eq. (37), but would suffer from angular wraparound issues at the −*π* → *π* boundary. To avoid this problem, we will adopt a 2 − *D* formalism.

### Softening of circular constraints and shift to Cartesian coordinates

To avoid angular wraparound issues, we consider the angular representation in *x, y* coordinates: *P* (*θ*_*t*_|*σ*_*t*_, *θ*_*t*−1_) = *P* (*x*_*t*_, *y*_*t*_|*σ*_*t*_, *x*_*t*−1_, *y*_*t*−1_). This will require us to soften the strict circular constraint that is intrinsic when using only *θ*.

We assume that the angular velocity is well approximated in Cartesian coordinates by the small angle approximation

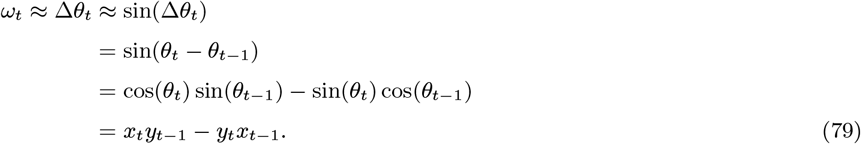

We consider a population of Poisson neurons responding to the angular velocity through a kernel *ϕ*_*t*_ = *ϕ*(*ω*_*t*_) ≈ *ϕ*(*x*_*t*_*y*_*t*−1_−*y*_*t*_*x*_*t*−1_) and a soft ring prior of radius *ρ* with form 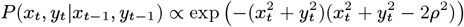 (Fig. 4B-C). These choices allow us to rewrite Eq. (73) as:

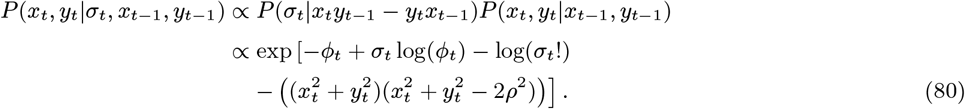

Taking the log, we get

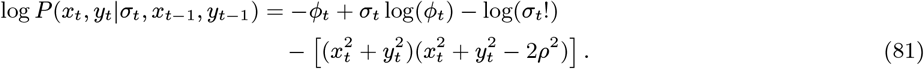

The exact form of the Langevin sampler will depend on the choice of the firing rate nonlinearity of the Poisson neurons. For example, using an exponential kernel

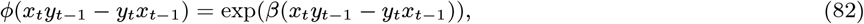

and taking the gradient ∇_*x,y*_ log *P*, leads to

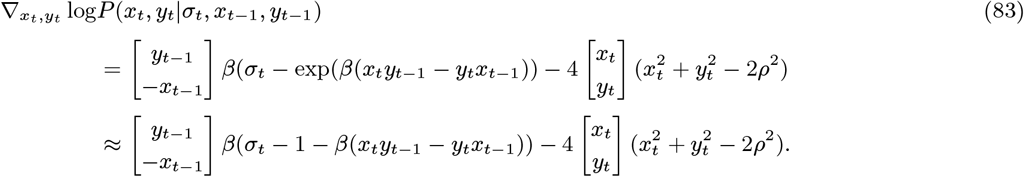

Considering the Langevin sampler in discrete time, and assuming that 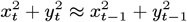, one gets

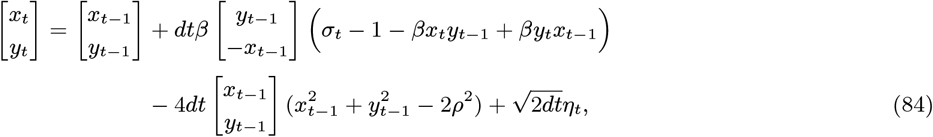

where *η* ∼ N (0, **I**). Considering now that

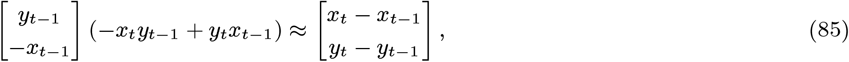

and moving the terms in ·_*t*_ to the left-hand side, simplifying, and considering *α* = 1 + *dtβ*^⊤^*β*,

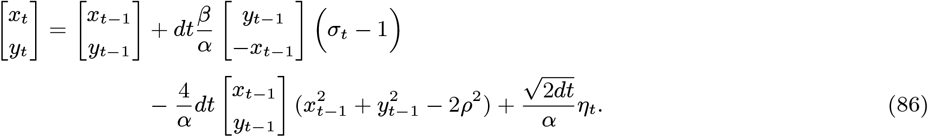

This equation resembles a noisy 2 − *D* angular velocity integrator informed by *σ*_*t*_, with a nonlinear part that implements a soft circular constraint, whose strength is regulated by *γ*, and takes the form of a limit cycle of radius *ρ* [115]. This general procedure could be extended to more complicated examples, which we will explore in future work. All in all, these models show how sampling dynamics and attractor dynamics can be combined in spiking neural networks.

Eq. (86) can be implemented into a SCN by using Eq. (37); In detail, for the simulations in Fig. 4 and Fig. 5, this is the voltage equation being used:

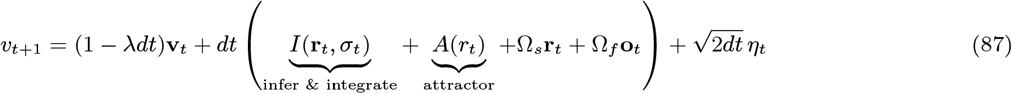

where

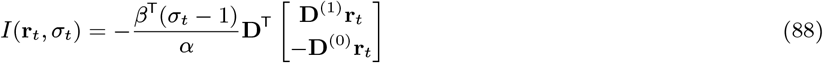

with **D**_(*i*)_ denoting the *i*−th row of **D**^⊤^, and

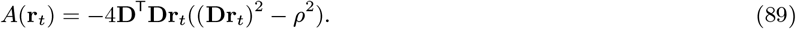

The other terms are defined as in Eq. (37); for completeness, **Ω**_*s*_ = **D**^⊤^**D, Ω**_*f*_ = −**D**_*t*_**D**, and *η* = **D**^⊤^*ξ* where *ξ* ∼ N (0, **I**).

### Multimodal integration and possible reset mechanisms

The HD systems receives multiple inputs. Moreover, under normal conditions, the HD system can reset towards a known landmark,; this necessitates the integration of visual and angular velocity cues. This process could be implemented in various ways. For example, there could be a mechanism that resets the network readout to a set value (e.g., 0) every time the fly sees the sun. Consider an external input to the network, *σ*^(*reset*)^, that interacts with the other terms as in

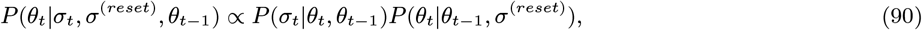

and the prior *P* (*θ*_*t*_|*θ*_*t*−1_, *σ*^(*reset*)^) is such that *P* (*θ*_*t*_|*θ*_*t*−1_, *σ*^(*reset*)^) = *P* (*θ*_*t*_|*θ*_*t*−1_) if *σ*^(*reset*)^ = 0, and *P* (*θ*_*t*_|*θ*_*t*−1_, *σ*^(*reset*)^) ∼ 𝒩 (0, *f* (*σ*^(*reset*)^)) otherwise.

A slightly more realistic example could entail an external input that encodes the direction of a known landmark, again at *θ* = 0, noisly, such as *P* (*σ*^(*reset*)^|*θ*) ∝ Poiss(*c* exp(−*θ*^2^)) and *P* (*θ*_*t*_|*θ*_*t*−1_, *σ*^(*reset*)^) = *P* (*σ*^(*reset*)^|*θ*_*t*_)*P* (*θ*_*t*_|*θ*_*t*−1_). In that case,

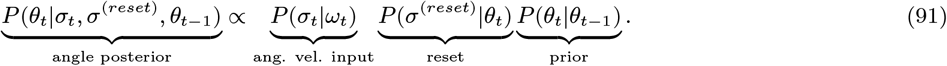

This can be implemented in 1 − *D* as

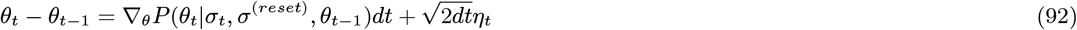

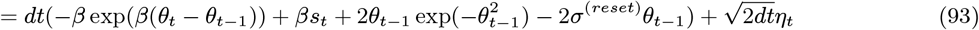

and in 2 − *D*, assuming *P* (*σ*^(*reset*)^|*x, y*) ∝ Poiss(*c* exp(−(*x* − 1)^2^ − *y*^2^)), as

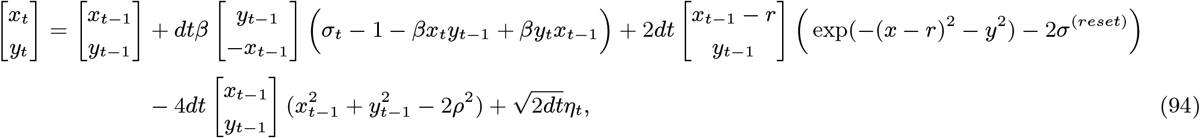

## Code availability and Github repository

The code to reproduce the analyses in the main figures is available at https://github.com/michnard/sampl_attract_HD.

## CRediT Author contributions

**Vojko Pjanovic:** Analysis, Investigation, Software, Validation, Visualization, Methodology, Writing

**Jacob Zavatone-Veth:** Conceptualization, Investigation, Methodology, Writing

**Paul Masset:** Conceptualization, Methodology

**Sander Keemink:** Conceptualization, Funding Acquisition, Investigation, Methodology, Project Administration, Supervision, Visualization, Writing

**Michele Nardin:** Conceptualization, Analysis, Funding Acquisition, Investigation, Methodology, Project Administration, Resources, Software, Supervision, Validation, Visualization, Writing

## Acknowledgments

We would like to express our gratitude to Ann Hermundstad and Brad Hulse for their invaluable insights and feedback on the project and the manuscript. We also acknowledge Shivam Chitnis, Farhad Pashakhanloo, Sandro Romani, and William Podlaski for their valuable feedback on early versions of this manuscript.

VP and MN are supported by the Howard Hughes Medical Institute. JAZV is supported by the Office of the Director of the National Institutes of Health under Award Number DP5OD037354. The content is solely the responsibility of the authors and does not necessarily represent the official views of the National Institutes of Health. JAZV is further supported by a Junior Fellowship from the Harvard Society of Fellows. This research was carried out in part thanks to funding from the Canada First Research Excellence Fund, awarded to PM through the Healthy Brains, Healthy Lives initiative at McGill University.

The cartoon of *Drosophila melanogaster* used in Figures 1, 3, and 4, was created in BioRender: https://BioRender.com/x68r143; the cartoon of a mouse used in Figure 1 was created in BioRender: https://BioRender.com/w64d461.

## Supplementary Figures

**Supp. Fig. 1.**
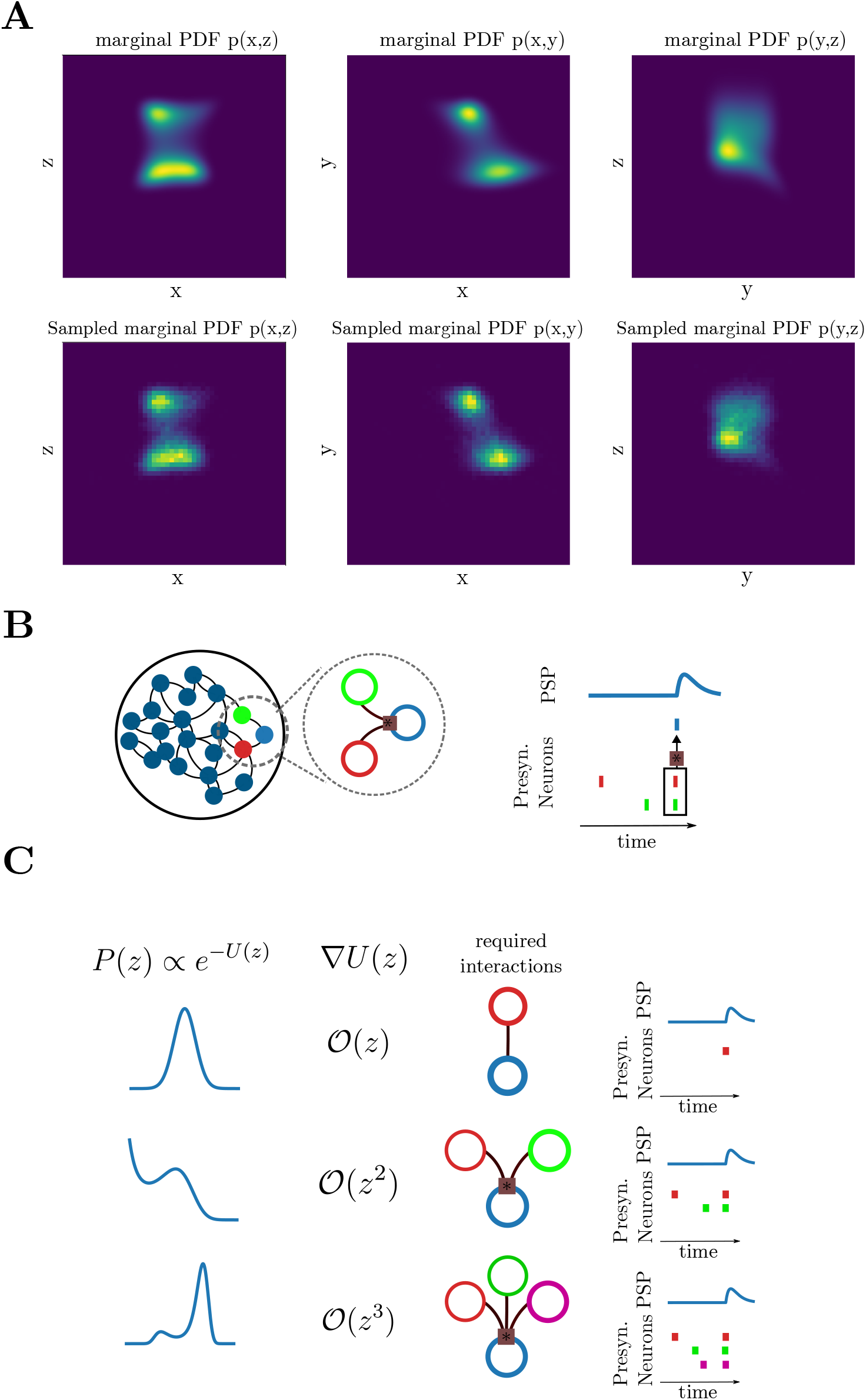
Sampling from high−*D* non-Gaussian distributions in mSCNs. **(A)** 3*D*, 4^*th*^ order polynomial moment exponential family distribution, given by exp(−*U* (*x, y, z*)), where *U* (*x, y, z*) = *x* + *x*^2^ − 3*x*^3^ + *y*^2^ − 3*y* + *x*^2^*y*^2^ + *xyz*^2^ + *x*^2^*yz* + *x*^4^ + *z*^4^. Top row: true marginal PDF, bottom: sampled marginal normalized histogram after 10000 samples. **(B)** Nonlinear polynomial dynamical systems in SCNs require multiplicative interactions among pairs of incoming inputs. The PSP of the receiving neuron *i* will be affected according to 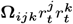, where *j, k* are the two presynaptic neurons, and **Ω** is the matrix of nonlinear synaptic weights. **(C)** Higher-order polynomials require increasingly complicated multiplicative interactions (Supp. Fig. 1B). Specifically, networks have to implement (*g* − 1)-th order multiplications for distributions with *g*-th order polynomial moments (see Methods).

**Supp. Fig. 2.**
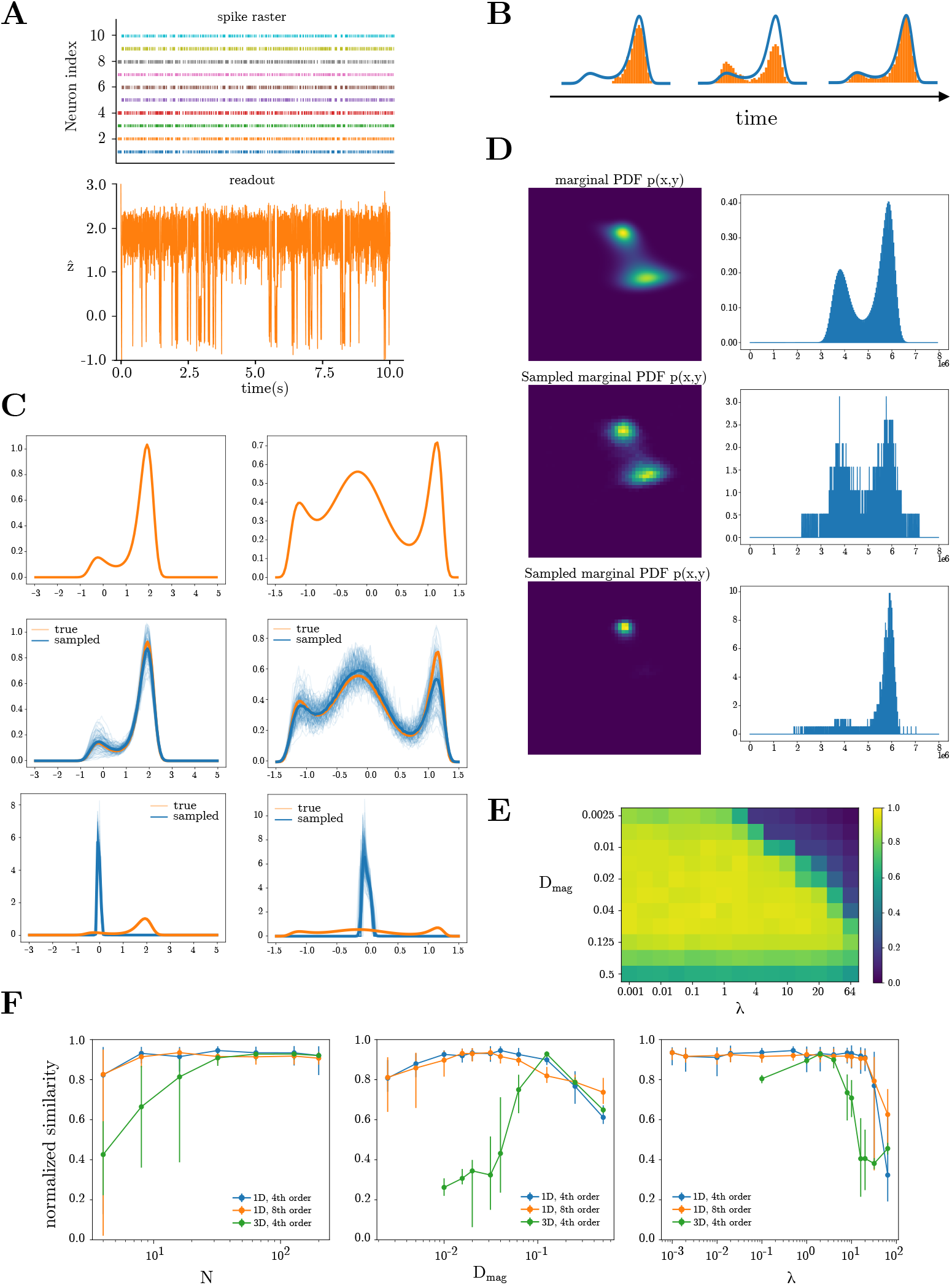
Sampling behaviour and performance for various distributions. **(A)** Example neural activity of 10 neurons (top) and linearly read out samples (bottom) for a bimodal distribution (as in panel C, left). **(B)** Schematic of the evolution over time of an empirical histogram-based density from samples (orange), and the true density (blue). **(C)** Visualization of good and bad sampling for the 1D, 4^th^ and 8^th^ order distributions (left and right, respectively); true distributions (top) are sampled with best (middle) and worst (bottom) parameters as per Table 2. Orange lines are the true distributions, thick blue line is the mean of 100 runs, each plotted as a thin blue line. **(D)** Visualization of good and bad sampling for the 3D, 4^th^ order distribution; true distribution (top) is sampled with the best and worst parameters, shown in middle and bottom respectively, as per Table 2. The left column showcases the marginal distributions *P* (*x, y*), and the right column contains the flattened 3D histogram. **(E)** Similarity scores as a function of *D*_*mag*_ and *λ* for the 1D, 4^th^ order distribution. **(F)** Similarity scores for characterizing the sampling performance for three distributions given by Eq. (96) with varying parameter sets. The optimal parameter set {*N, D*_*mag*_, *λ*} is determined for each distribution (Table 2), and consequently one parameter is varied while the other two are fixed. Blue, orange, and green line correspond to the 1D fourth order, 1D eighth order, and 3D fourth order distributions respectively. Each datapoint is the mean over 20 runs for that parameter set, and the corresponding bars span 5th to 95th percentiles. Similarity score is determined as per Perfomance calculation.

**Supp. Fig. 3.**
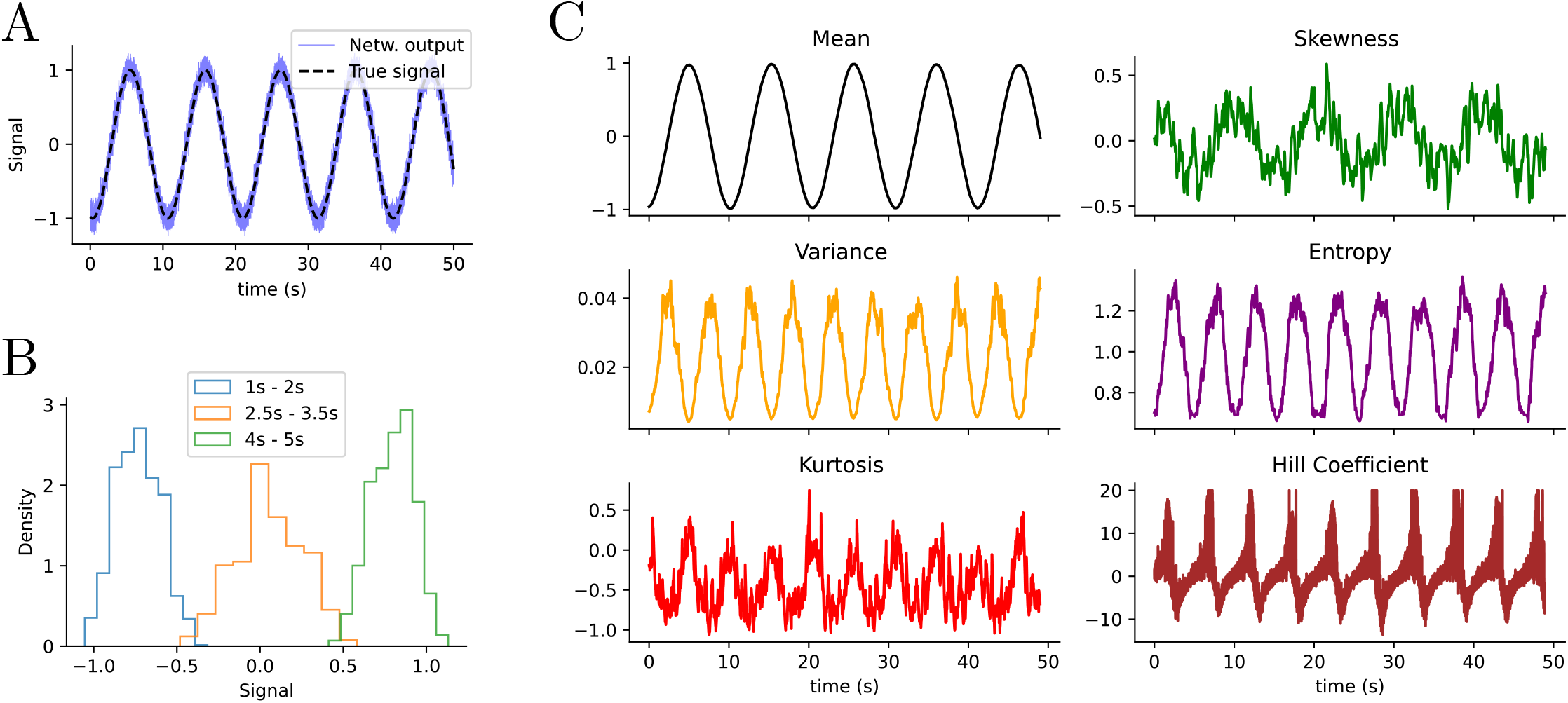
Sampling-based inference allows the computation of various statistics online on time-varying stimuli posterior distributions. An oscillating external stimulus is encoded by 100 Poisson neurons for 50 seconds with a *dt* = 10^−3^s. A SCN network with 50 neurons, following the dynamics of Eq. (53), computes real-time sampling-based inference. **(A)** True signal (dashed black) vs network output (light blue) over 50 seconds. **(B)** We assume that a downstream reader computes various statistics on a running window of 1000 samples. Three histograms show the real-time sampling process. **(C)** Various statistics measured on the running windows of 1000 samples: first 4 central moments, entropy measured on the discretized distribution, hill coefficient measured with a hill estimator with *k* = 100.

**Supp. Fig. 4.**
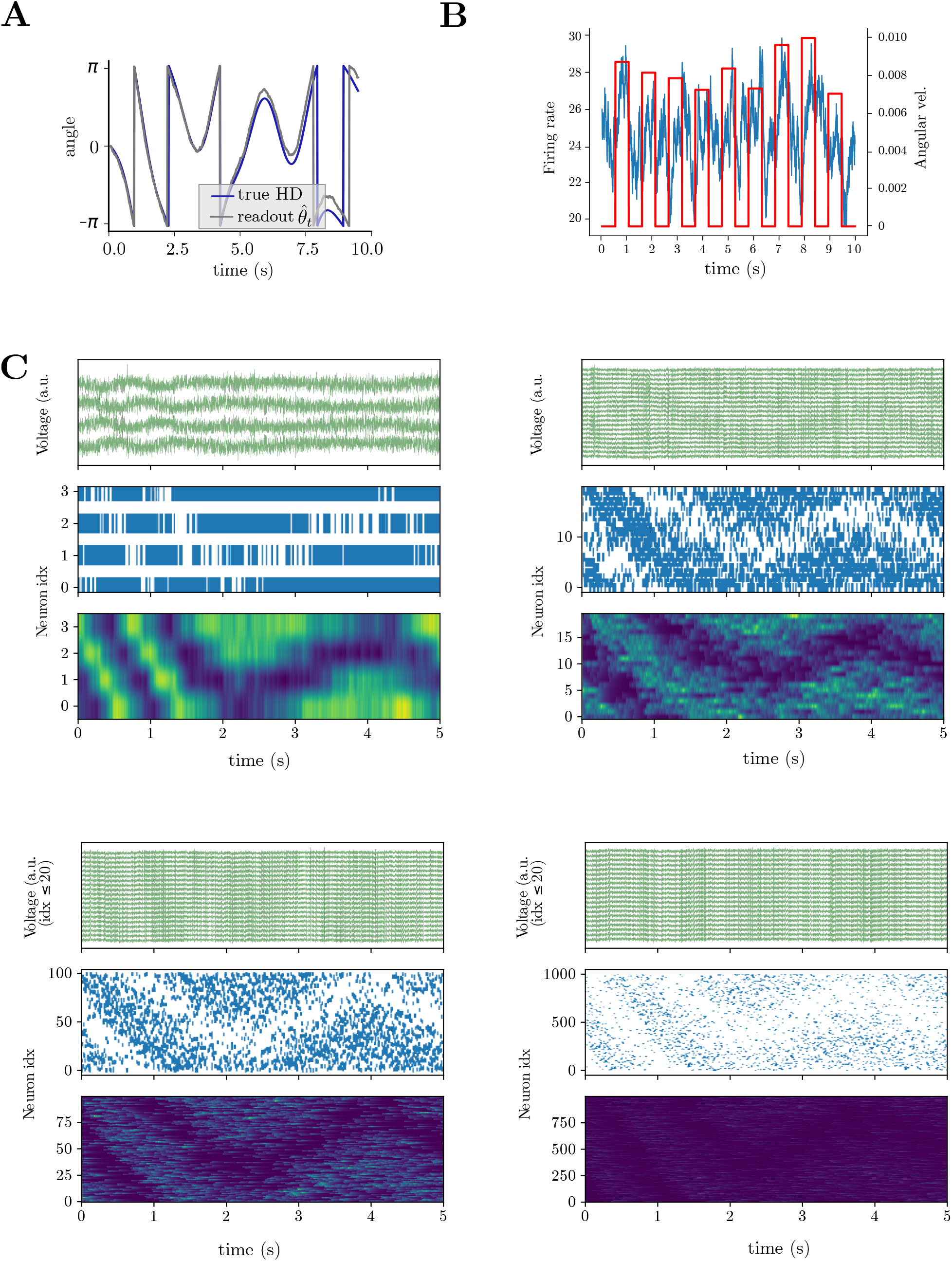
Biological phenomena captured by sampling-based HD network. **(A)** True HD vs SCN readout for an example network run as in Fig. 4. Notice that the network accumulates errors and loses the true orientation over time. **(B)** Average network activity (blue) during pulses of angular velocity (red). **(C)** Voltage (**v**), spikes (*o*), and filtered spikes (*r*) for SCN HD networks with 4, 16, 100, and 1000 neurons. Same inputs as in Fig. 4.

**Supp. Fig. 5.**
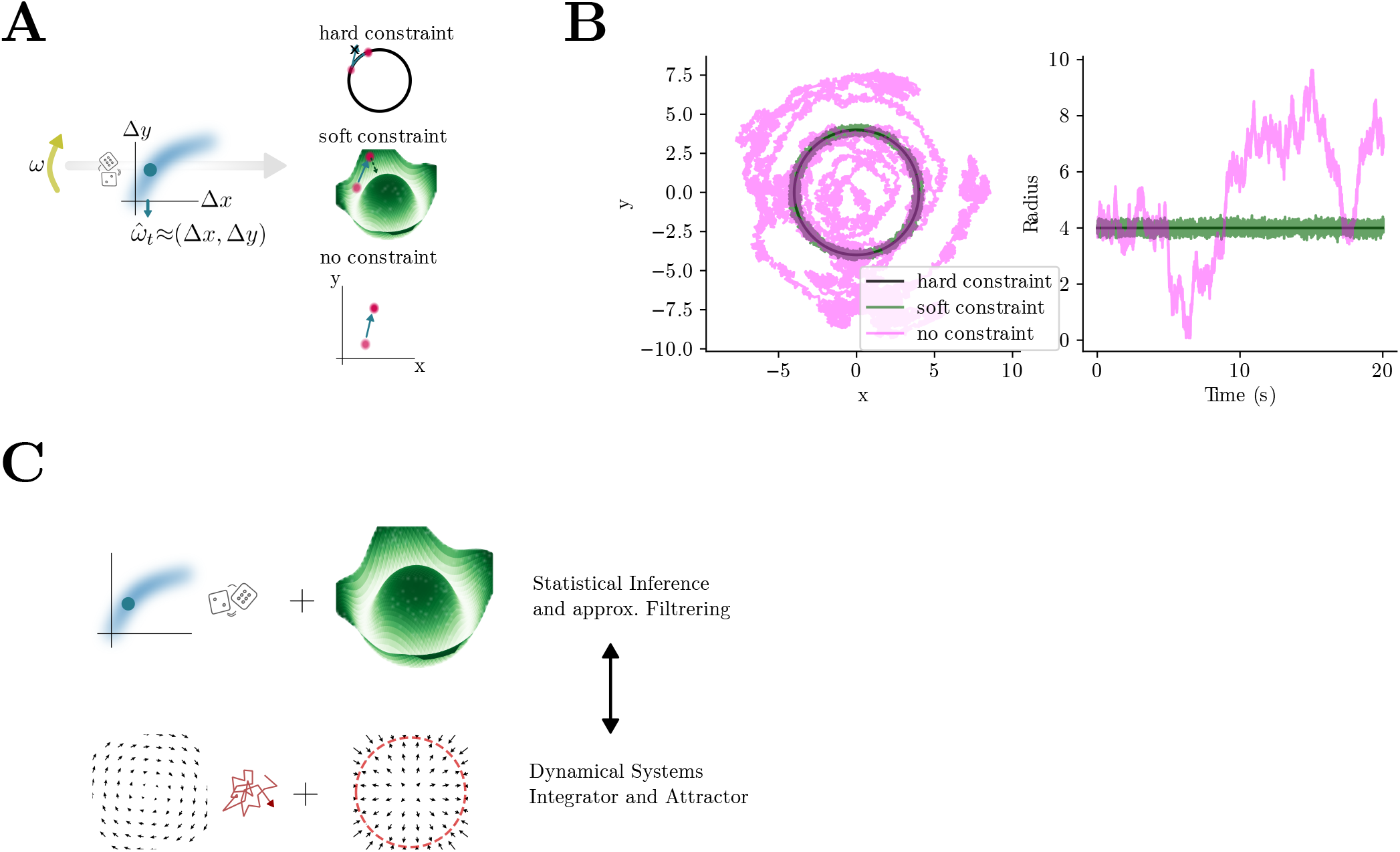
Constraining 2 − *D* angular representations. **(A)** Left: angular velocity *ω* is encoded noisily, and then transformed into a 2 − *D*, Δ*x*, Δ*y* HD update. Right: the HD representation in 2 − *D* can be constrained in various ways; top: a hard constrain represents a renormalization, whereby 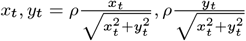; middle: a soft constraint biases the dynamics to return towards the circle by including an energy term away from the circle, as in (*x*^2^ + *y*^2^)(*x*^2^ + *y*^2^ − *ρ*^2^), so that the dynamics are driven towards the outside if *x*^2^ + *y*^2^ *< ρ*^2^, and towards the inside if *x*^2^ + *y*^2^ *> ρ*^2^; bottom: no constraints allow the dynamics to go wherever in the 2 − *D* plane. **(B)** Example *x, y* dynamics for hard (red), soft (green), and no constraints (cyan). Notice the radius oscillations (right panel). **(C)** Writing the approximate Baysian filtering problem through Langevin dynamics results into a nonlinear dynamical system resembling a 2 − *D* integrator combined with a circular limit cycle (i.e. attractor).

**Supp. Fig. 6.**
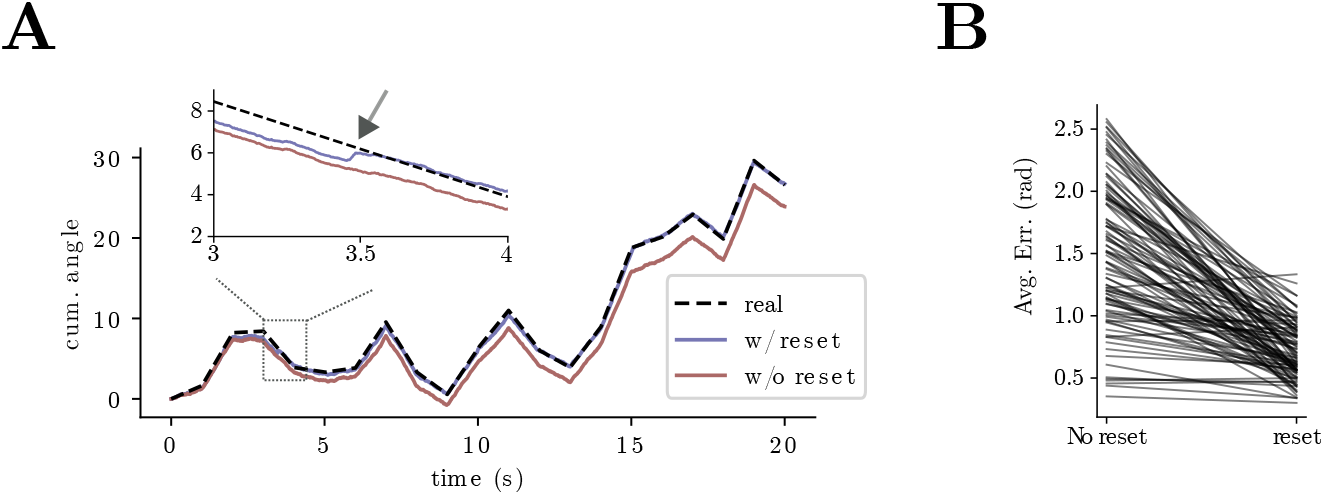
Implementing a visual reset mechanism through multimodal integration. **(A)** Comparison of cumulative angle for real vs. HD with reset vs. HD without reset over 20 seconds behavior. Inset: zoom in to show the effect of the reset. **(B)** Avg. error after 20 seconds behavior with vs. without reset mechanisms.

## Appendix

### Convergence analysis

To better understand the overall performance and the impact of the parameters on the sampling of the SCN, we chose three distributions and sampled from them using Eq. (37) with varying parameters:

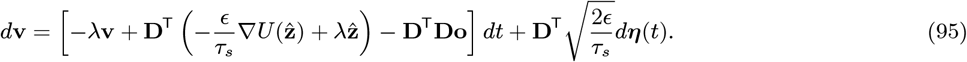

To appropriately test the sampling performance, we hand-picked three distributions that portrayed interesting and somewhat complicated characteristics: the simplest distribution is one-dimensional with fourth-order terms in the energy function, which results in a double-peaked distribution (Supp. Fig. 2C, left); the second distribution is also one-dimensional but with eighth-order terms in the energy function, leading to a triple-peaked distribution (Supp. Fig. 2C, right); lastly, we also analyzed a three-dimensional distribution with fourth-order terms in the energy function, as seen in (Supp. Fig. 1A). Specifically, the distributions are given by respectively

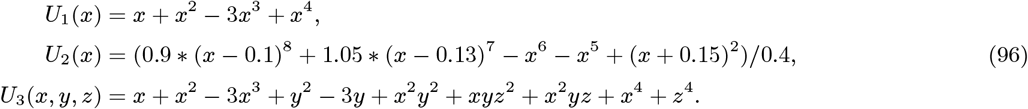

As for the parameters, we varied the number of neurons *N*, the voltage leak *λ*, and the magnitude of the decoding matrix *D*_*mag*_ = ∥**D**∥. The following parameter sets were used for sampling all distributions:

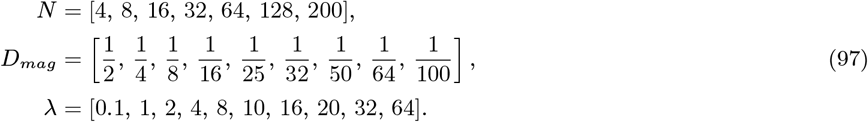

Additionally, for the 1D, 4^th^ and 8^th^ order distributions, 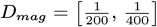 and 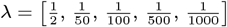 were added to the parameter sets for better characterization. We ran all permutations of the parameters to see which combination would produce the best result, with 20 runs for each permutation to make sure accurate results are obtained given the stochasticity in the sampling procedure. Other relevant parameters were standardized across all runs: for the 1D distributions, nt = 10000 timesteps, dt = 0.001 time step size, *ϵ* = 10 learning rate; for the 3D distribution, nt = 10000 timesteps, dt = 0.01 time step size, *ϵ* = 100 learning rate. No random seed was used for simulations, so the read-out matrices **D**, i.e. also the interaction matrices (Eq. (26)), as well as the noise *η* were randomized in every run.

#### Performance calculation

To quantify the performance of the sampling, we used the ℒ_1_ metric between the histogram bins of the ground truth distribution and the sampled empirical distribution with standardized bins,

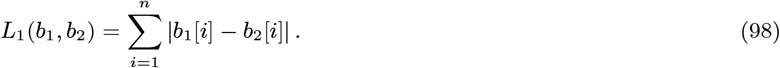

The histogram bins were standardized for each distribution to ensure validity: in the 1D 4th order case, 100 histogram bins between −3 and 5; in the 1D 8th order case, 100 bins between −1.5 and 1.5; in the 3D 4th order case, 200 bins between −4 and 4 in each dimension. It was ensured that the histograms of the ground truth distribution and empirical distribution had the same bin centers. The ground truth distribution and the empirical distributions were appropriately normalized: the ground truth distribution was normalized by dividing by the partition function through numerical integration over the entire space, and the empirical distribution was normalized using the density option in the `numpy.histogramdd` function.

Furthermore, as the ℒ_1_ metric magnitude varied strongly in each distribution ensemble, i.e. the collections of all sampling runs for a specific distribution in Eq. (96), we calculate a normalized similarity score for each distribution. Within each distribution ensemble, the best ℒ_1_ is taken as the performance baseline. Then, we compute the difference between each ℒ_1_ value and the best (minimum) ℒ_1_ in the dataset, effectively inverting the metric so that higher values correspond to better similarity. This ensures that a higher score represents a better match. To further normalize the scores for comparison across different distributions, we divide by the maximum value obtained after inversion, yielding a final similarity measure in the range [0, 1]. The transformed score is thus given by:

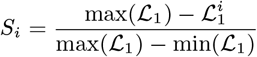

where *S*_*i*_ is the normalized similarity score for the *i*th distribution.

#### Results

The results of the simulations can be seen in Supp. Fig. 2. The sampled distributions for the best and worst parameter sets are visualized in Supp. Fig. 2C&D. To see the effects of each parameter on the sampling performance individually, we determined the optimal parameter sets {*N, D*_*mag*_, *λ*} (Table 2), and varied each parameter while the other two are fixed to the optimal value; the result is visualized in Supp. Fig. 2F.

**Table 2.** Best and worst parameter sets for sampling from the distributions given by Eq. (96), visualized in Supp. Fig. 2C&D.

| Distribution | Parameters |  |  |
| --- | --- | --- | --- |
| | $N$ | $D_{mag}$ | $\lambda$ |
| 1D, 4<sup>th</sup> order<br>best | 32 | $\frac{1}{25}$ | 0.5 |
| 1D, 4<sup>th</sup> order<br>worst | 64 | $\frac{1}{400}$ | 64 |
| 1D, 8<sup>th</sup> order<br>best | 16 | $\frac{1}{50}$ | $\frac{1}{100}$ |
| 1D, 8<sup>th</sup> order<br>worst | 32 | $\frac{1}{400}$ | 64 |
| 3D, 4<sup>th</sup> order<br>best | 128 | $\frac{1}{8}$ | 2 |
| 3D, 4<sup>th</sup> order<br>worst | 8 | $\frac{1}{50}$ | 1 |

Interpreting the result in Supp. Fig. 2F, we see that the number of neurons *N* does not influence performance in the 1D case, which is sensible from the geometric perspective [60], as 2 neurons with decoding vectors spanning positive and negative values suffice for optimal performance. For 3D distributions, the number of neurons impacts performance, due to the potential lack of closure in the bounding box for low *N* [55, 60]. Second, the magnitude of the read-out matrix **D** is most relevant to the sampling performance of the network, which shows a strong dependence on *D*_*mag*_. Both small and large magnitudes lead to bad results, for different reasons: small decoding vectors cannot counteract the decay, while big decoding vectors are too imprecise to obtain a reasonable readout [60]. Finally, the voltage decay *λ* has generally negligible impact on performance. Only when *λ >* 10 it affects the sampling negatively. This corresponds to a quick decay and, thus, a short memory of the voltage and readout. The network forgets the previous state and cannot properly sample the distribution. From Fig. 2E, we see that the sampling performance is not independently modulated by the parameters *D*_*mag*_ and *λ*; therefore, a potential solution for the posited problem is to increase *D*_*mag*_, as the spiking will have a larger effect on the readout and can counteract the quick decay [60], but this will generally decrease the precision of the readout.

## Footnotes

1 We abuse the notation and use *β* to denote the vector containing all weights associated with the Poisson input neurons.

